# Systematic characterization of *Drosophila* RhoGEF/GAP localizations uncovers regulators of mechanosensing and junction formation during epithelial cell division

**DOI:** 10.1101/2022.12.29.522184

**Authors:** Florencia di Pietro, Mariana Osswald, José M De las Heras, Ines Cristo, Jesus Lopez- Gay, Zhimin Wang, Stéphane Pelletier, Isabelle Gaugué, Adrien Leroy, Charlotte Martin, Eurico Morais-De-Sá, Yohanns Bellaïche

**Author notes:** co-first authors (Florencia di Pietro, Mariana Osswald). co-second authors (José M De las Heras, Ines Cristo).

## Abstract

Cell proliferation is central to epithelial tissue development, repair and homeostasis. During cell division, small RhoGTPases control both actomyosin dynamics and cell-cell junction remodelling to faithfully segregate the duplicated genome while maintaining tissue polarity and integrity. To decipher the mechanisms of RhoGTPases spatiotemporal regulation during epithelial cell division, we generated a transgenic fluorescently tagged library for *Drosophila* Rho Guanine exchange factors (GEF) and GTPase activating proteins (GAP), and systematically characterized their endogenous distributions by time-lapse microscopy. Thereby, we unveiled candidate regulators of the interplay between actomyosin and junctional dynamics during epithelial cell division. Building on these findings, we uncovered that during cytokinesis, Cysts and RhoGEF4 play sequential roles in mechanosensing and *de novo* junction formation, respectively. We foresee that the RhoGEF/GAP library will be a key resource to understand the broad range of biological processes regulated by RhoGTPases.

## INTRODUCTION

In animal cells, cell division entails drastic cell shape changes necessary for the faithful segregation of the duplicated genome into the two daughter cells. These morphological changes include cell rounding required for correct spindle formation and orientation, as well as cytokinesis to separate the daughter cell cytoplasms (Cadart et al., 2014; Glotzer, 2017). Studies in single cells and tissues have shown that these cell shape changes are powered by small RhoGTPases that remodel the actomyosin cytoskeleton (Cadart et al., 2014; Glotzer, 2017; Taubenberger et al., 2020). Moreover, in multicellular contexts, RhoGTPases are also critical to couple cell shape changes and junction dynamics to control tissue polarity, cohesion and architecture (Arnold et al., 2017; Buckley and St Johnston, 2022; Heasman and Ridley, 2008; Jaffe and Hall, 2005). Notably, cell division is tightly linked to cell fate specification as well as tissue growth, morphogenesis and mechanics (Godard and Heisenberg, 2019; Jülicher and Eaton, 2017; Lechler and Mapelli, 2021). Therefore, characterizing the regulation of RhoGTPases and its implication in cytoskeleton and junction dynamics during cell division in tissues is central to understand how cell number and genome integrity are controlled, and how tissue architecture and function are established and maintained.

The small RhoGTPases Rho, Rac and Cdc42 are key pleiotropic regulators of the actomyosin cytoskeleton and cell junction dynamics (Arnold et al., 2017; Buckley and St Johnston, 2022; Denk-Lobnig and Martin, 2019; Heasman and Ridley, 2008; Iden and Collard, 2008; Jaffe and Hall, 2005). They switch between an active GTP-bound state that binds downstream effectors, and an inactive GDP-bound state. These states are primarily regulated by Rho guanine exchange factors (RhoGEF) that activate RhoGTPases by exchanging GDP for GTP, and by RhoGTPase activating factors (RhoGAP) that promote GTP hydrolysis to GDP thereby inactivating RhoGTPases (Hodge and Ridley, 2016). Individual RhoGEF/GAP associated with the regulation of cell shape changes, cell division, migration and polarity have been identified in cultured cells (Arnold et al., 2017; Bagci et al., 2020; Jordan and Canman, 2012; Lawson and Ridley, 2018; Nakajima and Tanoue, 2011; Toret et al., 2014; Zihni et al., 2014), and to a lesser extent by targeted RNAi or mutant analyses in multicellular contexts (Fic et al., 2021; Garcia De Las Bayonas et al., 2019; Laurin et al., 2019; Mason et al., 2016; Silver et al., 2019; Toret et al., 2018). In addition, by ectopically expressing all human RhoGEF/GAP in cultured cell lines, a recent study has defined their localizations and biochemical interactomes, enabling a better understanding of single cell migration (Bagci et al., 2020; Müller et al., 2020). Therefore, the mechanisms mediating the spatiotemporal activation of small RhoGTPases are best understood in individual cells in interphase. However, the spatiotemporal regulations of RhoGTPases remain far less explored during cell division or in tissues, impeding our understanding of actomyosin and junction dynamics during interphase and cell division in epithelial tissues.

During animal cell cytokinesis, RhoGTPases control the pronounced cell deformations associated with cytokinetic ring constriction (Glotzer, 2017). Numerous studies have converged to show that the assembly and constriction of the actomyosin cytokinetic ring is powered by the membrane redistribution of the RhoGEF ECT2 (*Drosophila* Pebble, Pbl) and RacGAP1 *(Drosophila* Tumbleweed, Tum) within the dividing cell (Glotzer, 2017; Mishima et al., 2002; Su et al., 2011; Yüce et al., 2005; Zhao and Fang, 2005). Despite these fundamental findings, the role of most RhoGEF/GAP during cell division remains poorly explored. In addition, in epithelial tissues, several studies have shown that the drastic cytokinesis cell shape changes are coupled with E-Cadherin (Ecad) *adherens* junction (AJ) remodelling and *de novo* AJ formation (Herszterg et al., 2014; Ragkousi and Gibson, 2014). In particular, epithelial cytokinesis shares general features in several vertebrate and invertebrate tissues: i) *de novo* AJ formation is coordinated with cytokinesis, and relies on mechanosensing processes involving the dividing cell and its neighbours; and (ii) the arrangement of the newly-formed cell junctions is defined in late cytokinesis, and it is proposed to modulate tissue topology and morphogenesis (Firmino et al., 2016; Founounou et al., 2013; Gibson et al., 2006; Guillot and Lecuit, 2013; Herszterg et al., 2013, 2014; Higashi et al., 2016; Lau et al., 2015; McKinley et al., 2018; Morais-De-Sá and Sunkel, 2013; Pinheiro et al., 2017; Ragkousi and Gibson, 2014). Some of these epithelial cytokinetic features are known to be regulated by the Rho and Rac GTPases (Herszterg et al., 2013; Pinheiro et al., 2017), but the mechanisms controlling their local and temporal activations remain unknown.

Towards achieving an integrated view of the spatiotemporal regulation of actomyosin and junction dynamics in proliferative epithelia *in vivo*, we assembled a complete library of fluorescently tagged *Drosophila* RhoGEF/GAP. We then systematically analysed RhoGEF/GAP localizations from interphase to cell division in two *Drosophila* epithelial tissues by time-lapse microscopy. By doing so, we unravelled a series of putative regulators of epithelial tissue organization, polarity and dynamics. These results led us to focus on the RhoGEF Cysts and RhoGEF4, and to characterize their respective roles in mechanosensation and AJ formation during epithelial cell division. Altogether, our work advances the understanding of cell division in epithelial tissues, and highlights that the RhoGEF/GAP library will be a relevant resource to investigate how actomyosin and junction dynamics are controlled during development, repair and homeostatic processes.

## RESULTS

### Generation of a library of tagged RhoGEF/GAP and characterization of their localizations in interphasic epithelial cells

Despite the critical roles of RhoGTPases in all epithelial tissues, a systematic characterization of all RhoGEF/GAP localizations has not been performed in any tissue *in vivo*. To enable the systematic exploration of RhoGEF/GAP localizations, we assembled a library of *Drosophila* fluorescently tagged RhoGEF/GAP (26 RhoGEF, 22 RhoGAP, see Table S1 for tag positions and human orthologues). We first generated Bacterial Artificial Chromosomes (BAC) transgenic lines for 9 GFP-tagged RhoGEF/GAP. The advent of CRISPR/Cas9 methods led us to switch to tagging by CRISPR/Cas9 mediated homologous recombination. We thus produced GFP- or mKate2-tagged insertions for the remaining 39 RhoGEF/GAP (Fig. S1A). Among the generated CRISPR-Cas9 tagged lines, 34 out of the 39 (∼87%) tagged alleles were homozygous viable. We also built a plasmid library of homologous recombination donors and guide RNA expressing vectors to readily edit each RhoGEF/GAP locus to create loss of function alleles, and then facilitate structure-function analyses, or add any others tags for live-imaging or biochemical studies (Fig. S1B and Table S1).

As most of *Drosophila* RhoGEF/GAP localizations have not been yet assessed in polarized epithelial cells *in vivo,* we performed a detailed characterization of all RhoGEF/GAP localizations in interphase in epithelial tissue (Fig.S1, Fig. S2, Table S2 and not shown). This was performed in two distinct tissues characterized by different structural organization and mechanical properties: the pupal dorsal thorax epithelium of the pupa (notum) (Fig. S1C), and the follicular epithelium (FE) of the adult ovary during their proliferative stages (Fig. S1D). First, this systematic assessment of 48 RhoGEF/GAP confirmed the interphase localization of Cysts (Fig. S1E, S1F), RhoGEF2 (Fig. S1E, S1F), RhoGAP71E (Fig. S1E), Spg (Fig. S1E), Conu (Fig. S1E, S1F), RhoGAP19D (Fig. S1G, S1H), RtGEF (Fig. S1I), Pbl (Fig. S1J) and Tum (Fig S1J, S1K) observed in diverse *Drosophila* epithelial tissues (Dent et al., 2019; Fic et al., 2021; Garcia De Las Bayonas et al., 2019; Jones et al., 2010; Mason et al., 2016; Neisch et al., 2013; Prokopenko et al., 2000; Schmidt et al., 2018; Silver et al., 2019; Toret et al., 2018).

Second, our screen uncovered numerous additional RhoGEF/GAP localizations in each tissue (Fig. S2A, B and Table S2). Interestingly, these localizations could be associated with (i) the regulation of apico-basal polarity or actomyosin dynamics at the junctional, medio-apical or basal cortex, as exemplified by the distinct localizations observed for Graf (Fig. S2C and S2D), CdGAPr (Fig. S2C and S2G), RhoGAP1A (Fig. S2C and S2G), GEFMeso (Fig. S2E), Sos (Fig. S2E), Ziz (Fig. S2D), Zir (Fig. S2D), CG46491 (Fig. S2D), RhoGAP15B (Fig. S2G), Cdep (Fig. S2F and S2H) and RhoGAP5A (Fig. S2H); (ii) vertex organization or cell rearrangements (Mbc, RhoGAP5A, Cdep) (Fig. S2F and S2H-S2J); and (iii) nucleus related functions (RhoGAP54D, CG43102, CdGAPr and RhoGEF4) (Fig. S2K and S2L). Together, this initial characterization validated the fluorescently tagged library to explore the localizations of RhoGEF/GAP. We thus next explored the localizations of RhoGEF/GAP during epithelial cell division.

### Deciphering RhoGEF/GAP distributions during epithelial cell division

Epithelial cell division is characterized by a set of cell shape changes necessary to segregate the duplicated genome in the two daughter cells, and by *de novo* AJ formation needed for the maintenance of epithelial organization (Fig. 1A, (Glotzer, 2017; Herszterg et al., 2014; Ragkousi and Gibson, 2014; Taubenberger et al., 2020). Analysing MyoII dynamics is instrumental to investigate both cell shape changes and AJ dynamics during cell division (Herszterg et al., 2014; Ragkousi and Gibson, 2014; Ramkumar and Baum, 2016). We therefore systematically analysed the localization of each RhoGEF/GAP in conjunction with MyoII:3xmKate2 during cell division in the notum and the FE. This analysis confirmed the well-known localization of Pbl and Tum during mitosis and cytokinesis (Fig. S3A and S3B) (Mishima et al., 2002; Su et al., 2011; Yüce et al., 2005; Zhao and Fang, 2005). It further revealed a complex choreography of RhoGEF/GAP dynamics from prophase to cytokinesis and *de novo* junction formation (Fig. 1B and Table S2), including sets of RhoGEF/GAP accumulating around the nuclear envelope during prophase, labelling the spindle or the centrosomes during cell division (Fig. 1B, S3C, S3D and S3E). Here, we specifically describe cortical and junctional RhoGEF/GAP, for which the subcellular distributions could be associated with cell shape changes or *de novo* junction formation during cell division (Fig1C-J, FigS3F-N, Movies S1 and S2).

**Figure 1:**
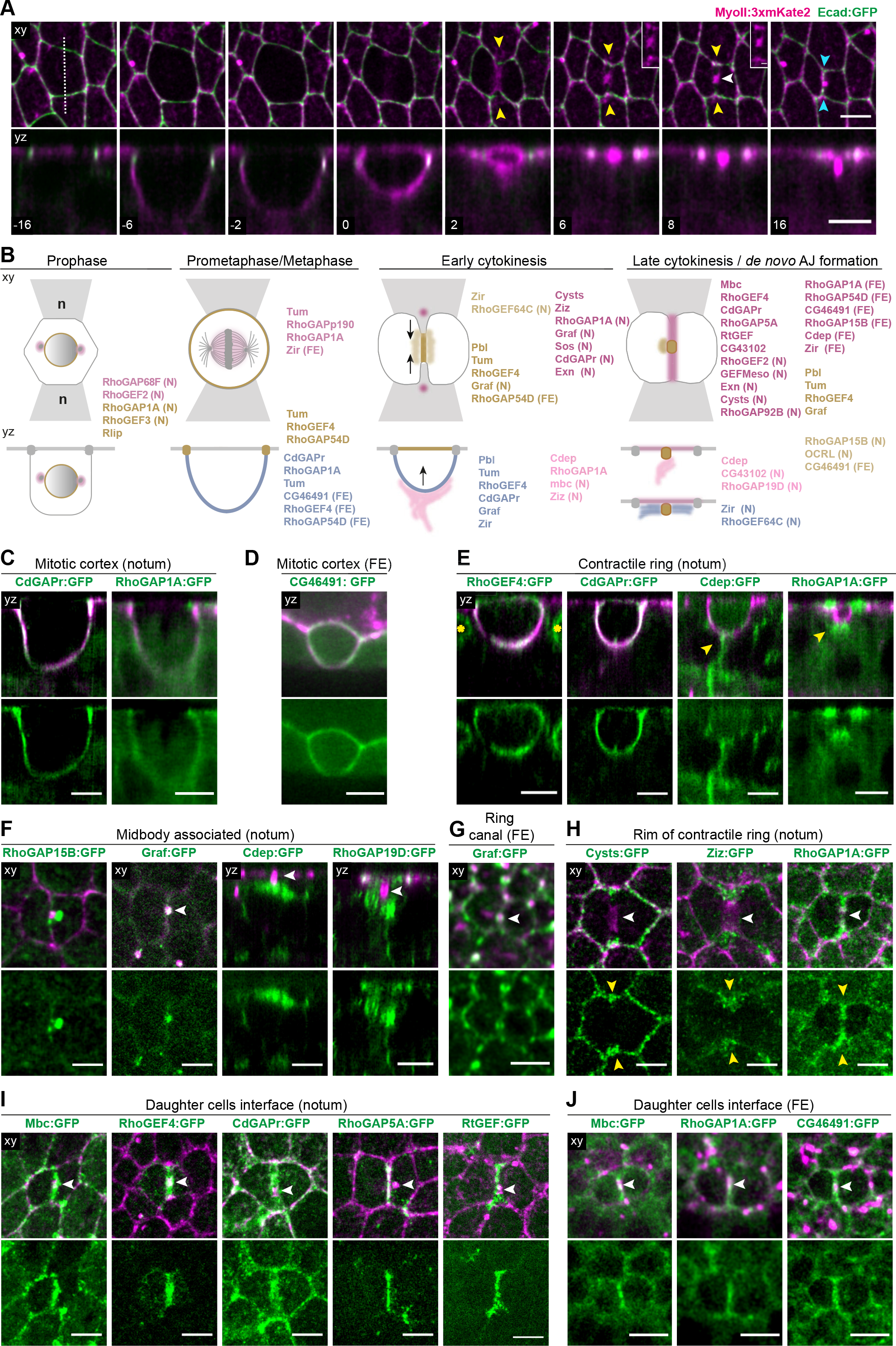
Localizations of RhoGEF/GAP during epithelial cell division. For all images, ‘xy’ indicates apical confocal top view at the level of the AJ in the notum and follicular epithelium (FE), while ‘yz’ denotes apical-basal confocal section at the level of the cytokinetic ring or the midbody for the notum, or a midsagittal section for the egg chamber. MyoII:3xmKate2 is shown in magenta. (A) xy (top) and yz (bottom) time-lapse images of Ecad:GFP and MyoII:3xmKate2 during cell division in the notum. Time (min) is set to 0 at the onset of cytokinesis marked by the initial deformation of AJ by the constriction of the cytokinetic ring. Cytokinetic furrowing occurs asymmetrically positioning the midbody apically (from t = 0 to t = 6 min). Upon midbody formation (t =6 min), a new AJ is formed between the two daughter cells (from t = 6 to t =16 min). Yellow arrowheads: MyoII accumulation in the neighbouring cells during ring constriction. White arrowheads: midbody. Cyan arrowheads: new daughter-daughter AJ. Inset: Close-up on the MyoII signal at the level of the future daughter cell interface. (B) Schematics of the subcellular localizations of selected RhoGEF/GAP during epithelial cell division in the notum and the FE in xy (top) and yz (bottom) views. The dividing cell is in white and the neighbouring cells (n) are in grey. The RhoGEP/GAP names are color-coded according to their localizations in each schematic of the different division phases. Black arrows: direction of cytokinetic furrowing. (N) and (FE) indicate localizations exclusively observed in the notum or the FE, respectively. Only RhoGEF/GAP with a good signal to noise ratio are indicated. See also Table S2 for a complete description. (C) yz images of CdGAPr:GFP, RhoGAP1A:GFP (top and bottom) and MyoII:3xmKate2 (top) in metaphase cells in the notum. (D) yz images of CG46491:GFP (top and bottom) and MyoII:3xmKate2 (top) in mitotic cells prior to anaphase in the FE. (E) yz images of RhoGEF4:GFP, CdGAPr:GFP, Cdep:GFP, RhoGAP1A:GFP (top and bottom) and MyoII:3xmKate2 (top) during cytokinetic ring contraction in the notum. RhoGEF4:GFP is localized a the nuclear envelope membrane in neighbouring interphasic cells (yellow asterisks, see also Fig. S5A). Yellow arrowheads: Cdep:GFP and RhoGAP1A:GFP enrichments below the cytokinetic ring. (F) xy images of RhoGAP15B:GFP, Graf:GFP (top and bottom) and MyoII:3mKate2 (top), and yz images of Cdep:GFP, RhoGAP19D:GFP (top and bottom), and MyoII:3xmKate2 (top) during late cytokinesis in the notum. White arrowheads: midbody. (G) xy images of Graf:GFP (top and bottom) and MyoII:3xmKate2 (top) during ring canal formation in the FE. White arrowhead: ring canal. (H) xy images of Cysts:GFP, Ziz:GFP, RhoGAP1A:GFP (top and bottom) and MyoII:3xmKate2 (top) during cytokinetic ring constriction in the notum. White arrowheads: cytokinetic ring. Yellow arrowheads: Cysts:GFP, Ziz:GFP and RhoGAP1A:GFP enrichments in the vicinity or at the rim of cytokinetic ring. (I) xy images of Mbc:GFP, RhoGEF4:GFP, CdGAPr:GFP, RhoGAP5A:GFP, RtGEF:GFP (top and bottom) and MyoII:3xmKate2 (top) during late cytokinesis or *de novo* daughter cells junction formation in the notum. White arrowheads: midbody. (J) xy images of Mbc:GFP, RhoGAP1A:GFP, CG46491:GFP (top and bottom) and MyoII:3xmKate2 (top) during late cytokinesis or *de novo* daughter cells junction formation in the FE. White arrowheads: midbody. Scale bars: 5µm (A, C, D, E, F, G, H, I, J), 1µm (inset in A). See also Figure S1-S3 and Movies S1 and S2.

#### RhoGEF/GAP localizations associated with cell shape changes during mitosis

Entry into mitosis is accompanied by cell rounding and a reorganization of actomyosin and junction complexes along the apical-basal axis of epithelial cells (Aguilar-Aragon et al., 2020; Ramkumar and Baum, 2016; Rosa et al., 2015; Taubenberger et al., 2020). We found that mitotic entry concurs with significant redistributions of several RhoGEF and GAP. In particular, we observed that RhoGEF4 and RhoGAP54D redistributed from the nucleus to the cortex in both tissues (Fig. S3F). In parallel, and mirroring the dynamics of MyoII (Fig. 1A), several RhoGEF/GAP expand along the basolateral cortex. This includes CdGAPr and RhoGAP1A in both tissues (Fig. 1C) and CG46491 in the FE (Fig.1D). During cell elongation in anaphase, a subset of RhoGEF/GAP were enriched or depleted from the cell poles in the FE: Mbc accumulated at the cell poles, while the RhoGEF CG46491 was cleared from the cell poles (Fig. S3G, Movie S2A and S2C). Together, this set of RhoGEF/GAP are good candidates to regulate the actomyosin cytoskeleton or cell junctions during the highly conserved processes of mitotic rounding, anaphase elongation and polar relaxation.

#### RhoGEF/GAP localizations associated with ring constriction and midbody formation

In addition to Pbl and Tum which are known to localize at the cytokinetic ring and the midbody (Glotzer, 2017; Mishima et al., 2002; Su et al., 2011; Yüce et al., 2005; Zhao and Fang, 2005), we identified a large set of RhoGEF/GAP with distinct localizations during cytokinesis (Fig. 1E-1J, and S3H-S3N). During early cytokinesis, RhoGEF4 (Fig. 1E and S3I), Graf (Fig. S3H), CdGAPr (Fig. 1E and S3I) and Zir (Fig. S3H) localized around the contractile ring. Interestingly, in the notum, we detected several RhoGEF/GAP localized in a position basal to the cytokinetic ring in previously uncharacterized filamentous or membranous structures. This includes Cdep (Fig. 1E, Movie S1A), RhoGAP1A (Fig. 1E), Mbc (Fig. S3H) and Ziz (Fig. S3H). Their localization basal to the ring was observed at different timepoints during ring contraction suggesting distinct functions. Upon completion of ring constriction, another subset of RhoGEF/GAP was observed associated to the midbody (Fig. 1F and Fig. S3J), localizing either transiently RhoGAP15B (Fig. 1F), RhoGEF4 (Fig. 1I), OCRL (Fig. S3J), CG46491 and Mbc (Fig. 1J) or more stably (Graf, Fig. 1F) around or at the midbody. Furthermore, we found specific localizations that might be associated with the distinctive midbody dynamics or functions known in each tissue. In the notum, as the midbody moves basally during septate junction formation and abscission (Daniel et al., 2018; Wang et al., 2018), we found that Cdep (Fig. 1F, Movie S1A), RhoGAP19D (Fig. 1F) and CG43102 (Fig. S3J) accumulated underneath the midbody. In the FE, where abscission is arrested to form ring canals (Airoldi et al., 2011; Chaigne and Brunet, 2022), Graf was initially enriched at the midbody and then accumulated on each side of the newly formed ring canal (Fig. 1G, Movie S2B). Altogether, this group of RhoGEF/GAP provides entry points to further understand basic features of epithelial division, such as asymmetric furrowing, midbody dynamics, junction formation or abscission regulation.

#### RhoGEF/GAP localizations associated with de novo AJ formation

At least two distinct processes couple cytokinesis and *de novo* AJ formation in epithelial tissues (Founounou et al., 2013; Herszterg et al., 2013; Higashi et al., 2016; Morais-De-Sá and Sunkel, 2013; Pinheiro et al., 2017). The first one is linked to cytokinetic ring contraction, while the second one occurs upon midbody formation. During ring constriction, the dividing and neighbouring cells membranes co-ingress, and the contractile force of the ring triggers a mechanosensing process leading to junctional or actomyosin reorganization at the rim of apical cytokinetic ring in the neighbouring cells (Herszterg et al., 2013; Higashi et al., 2016; Pinheiro et al., 2017). Upon midbody formation, the second process involves a Rac- and Arp2/3-dependent F-Actin accumulation at the midbody and at the prospective daughter cell interface. This promotes the withdrawal of the ingressed neighbouring cell membranes and *de novo* daughter-daughter AJ formation (Herszterg et al., 2013; Morais-De-Sá and Sunkel, 2013). We therefore examined RhoGEF/GAP distributions at the rim of the apical cytokinetic ring during its constriction, and at the level of the prospective AJ upon midbody formation. Notably, in one or both tissues, Cysts (Fig. 1H), Ziz (Fig. 1H, Movie S1I), RhoGAP1A (Fig. 1H, Movie S1F), Graf (Fig. S3J, Movie S1D), CdGAPr (Fig. S3K, Movie S1B) and Exn (Fig. S3K, Movie S1C) can be found at the rim of cytokinetic ring, where MyoII accumulates (Fig1A, (Herszterg et al., 2013)). Furthermore, we found specific RhoGEF/GAP accumulating transiently near or at the daughter cell interface during cytokinesis and *de novo* junction formation. These include Mbc (Fig. 1I and 1J, Movies S1E and S2C), RhoGEF4 (Fig. 1I), CdGAPr (Fig. 1I, Fig. S3N, Movie S1B), RhoGAP5A (Fig. 1I, Fig. S3N, Movies S1G and S2E) and RtGEF (Fig. 1I, Movie S1H), both in the notum and FE, CG43102 (Fig. S3M), Exn (Fig. S3M, Movie S1C) and GEFMeso (Fig. S3M) in the notum, as well as RhoGAP1A (Fig. 1J, Movie S2D), CG46491 (Fig. 1J, Movie S2A), Cdep (Fig. S3L), RhoGAP15B (Fig. S3N, Movie S2F), Zir (Fig. S3N) and RhoGAP54D (Fig. S3N) in the FE.

Collectively, the family-wide RhoGEF/GAP localization analyses revealed a rich spatiotemporal pattern of RhoGTPases regulators in proliferative epithelial tissues. This provides a large set of candidate regulators of cell division and junction remodelling in the polarized and multicellular context of epithelial tissues. In addition, the comparative analyses of RhoGEF/GAP localizations in two epithelial tissues are relevant to pinpoint: (i) RhoGEF/GAP harbouring similar localizations, evocative of general roles in epithelial tissues (e.g. Graf, Cysts, CdGAPr or RhoGEF4, Fig.1B and Table S2); (ii) RhoGEF/GAP exhibiting distinct distributions, advocating for tissue-specific activities associated with the different functions and dynamics of the two epithelial tissues (e.g. RhoGAP5A, RtGEF, Cdep and CG46491, Fig. 1B and Table S2). Next, to decipher the mechanisms of epithelial cytokinesis, we decided to focus on Cysts and RhoGEF4, which showed local enrichments at different phases of junction remodelling in both tissues (Fig. 1B, 1 H, 1I, Table S2). Using the aforementioned CRISPR/Cas9 plasmid library for genome editing of each RhoGEF/GAP, we generated and validated *cysts* and *rhogef4* null alleles to investigate their functions, mainly focusing on the notum epithelium.

### The RhoGEF Cysts plays a role in mechanosensing and daughter cell membrane juxtaposition during cytokinesis

Cytokinetic ring contraction triggers a mechanosensing response in the neighbouring cells associated with junctional remodelling during cytokinesis (Higashi et al., 2016; Pinheiro et al., 2017). As defined in the notum epithelium, ring contraction induces self-organized Rok and MyoII flows and accumulations at the rim of the cytokinetic ring in the neighbouring cells, thereby promoting the juxtaposition of daughter cell membranes (Herszterg et al., 2013; Pinheiro et al., 2017). During ring contraction, the Rho1 GTPase is also necessary to enhance MyoII accumulation in the neighbouring cells (Pinheiro et al., 2017). Having uncovered that Cysts, a known GEF for the Rho1 GTPase in *Drosophila* (Silver et al, 2019) is localized at the rim of the cytokinetic ring (Fig 2A, FigS4A, Movie S3A), we decided to investigate its function in mechanosensing during cytokinesis. Interestingly, the vertebrate Cysts orthologue, p114RhoGEF, has recently been proposed to mediate mechanosensing in cultured cells (Acharya et al., 2018; Duszyc et al., 2021), but whether Cysts or p114RhoGEF function in mechanical responses *in vivo* or during cell division is unknown.

**Figure 2:**
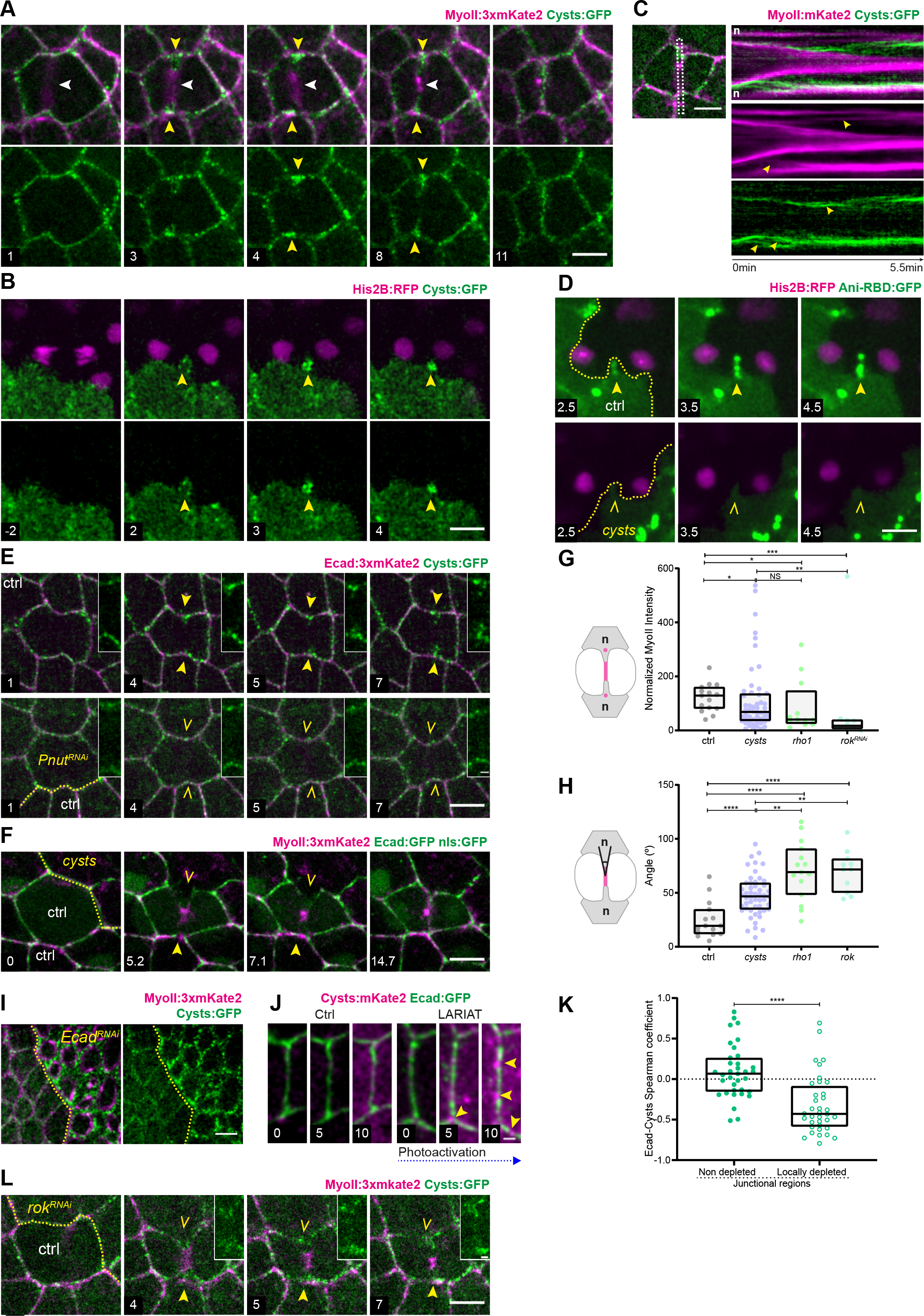
The RhoGEF Cysts modulates the mechanosensing response in the neighbouring cells, and daughter cells membranes juxtaposition. All images are apical top views at the level of the AJ in the notum. Unless otherwise specified, time (min, indicated in the lower left image corner) is set to 0 at the onset of cytokinesis marked by the initial deformation of AJ by the constriction of the cytokinetic ring. (A) Time-lapse images of Cysts:GFP (top and bottom) and MyoII:3xmKate2 (top) in the dividing cell and its neighbours. White arrowheads: cytokinetic ring. Yellow arrowheads: Cysts:GFP and MyoII:3xmKate2 accumulations at the rim of the cytokinetic ring. (B) Time-lapse images of clonally expressed His2B:RFP in the dividing cell (top) and Cysts:GFP (top and bottom) in its neighbours. Images are projections of the apical surface for the GFP channel, to highlight Cysts apical localization in the neighbour, merged with a projection from the apical cell surface to the nucleus for the RFP channel to visualize the dividing cell nuclei. Yellow arrowheads: Cysts:GFP accumulation within the ingressing membranes. (C) Apical view of Cysts:GFP and MyoII:3xmKate2 during epithelial cell division (left) and kymograph (right) of Cysts:GFP and MyoII:3xmKate2 within the region outlined in the apical view. Time (min) is set to 0 at the start of the movie. Yellow arrowheads. MyoII:3xmKate2 and Cysts:GFP speckles that flow between the ingressing membranes. n: neighbouring cells (D) Time-lapse images of Ani-RBD:GFP and His2B:RFP in control (ctrl) and *cysts* cells neighbouring a dividing cell. Ani-RBD:GFP and His2B:RFP are expressed clonally; *cysts* cells are marked by the loss of His2B:RFP signal in the bottom panels. A yellow dashed line marks the boundary between Ani-RBD:GFP expressing and non-expressing cells. Ani-RBD:GFP was analysed in cells adjacent to a cell that divides roughly parallel to them, and devoid of Ani- RBD:GFP signal. Ani-RBD:GFP accumulation was observed in 18 out of 22 ctrl neighbouring cells (81%), and in 5 out of 25 (20%) *cysts* neighbouring cells. Yellow arrowheads: Ani- RBD:GFP at the rim of the cytokinetic ring in ctrl neighbouring cell. Yellow open arrowheads: absence of Ani-RBD:GFP accumulation in a *cysts* neighbouring cell. (E) Time-lapse images of Ecad:3xmKate2 and Cysts:GFP in the context of ctrl (top) and *pnut^RNAi^* (bottom) dividing cell. *pnut^RNAi^* cells are marked by the expression of CAAX:tBFP (not shown) and a yellow dashed line marks the boundary between ctrl and *pnut^RNAi^* cells. Yellow arrowheads: Cysts:GFP at the rim of the cytokinetic ring in neighbours of a ctrl dividing cell. Yellow open arrowheads: reduced Cysts:GFP accumulation in cells neighbouring a *pnut^RNAi^* dividing cell. Insets: close-ups on Cysts:GFP signal at the rim of the cytokinetic ring in ctrl and *pnut^RNAi^* cells. Cysts:GFP accumulation at the rim of cytokinetic ring during early cytokinesis was observed in 82% of cells neighbouring a ctrl dividing cell (n=29) and in 36% of cells neighbouring *pnut^RNAi^* dividing cells (n=22). (F) Time-lapse images of Ecad:GFP and MyoII:3xmKate2 in the context of a dividing cell surrounded by ctrl and *cysts* neighbouring cells. *cysts* cells are marked by the loss of nls:GFP signal, and a yellow dashed line marks the boundary between ctrl and *cysts* cells. Note that nls:GFP is not visible in all ctrl cells since their nuclei can be located more basally. Yellow arrowheads: MyoII:3xmKate2 accumulation and daughter-daughter interface juxtaposition in the ctrl neighbouring cell. Yellow open arrowheads: reduced MyoII:3xmKate2 accumulation and delayed daughter-daughter interface juxtaposition in *cysts* neighbouring cell context. (G) Schematic of MyoII neighbour accumulation (left) and box plot of MyoII accumulation (median ± interquartile range, right) at the rim of the contractile ring at 80% of apical ring contraction in ctrl, *cysts*, *rho1 and rok^RNAi^* cells neighbouring a dividing cell. Note that for the *cysts* quantification, both the dividing and the neighbouring cells can be *cysts* mutant cells. (H) Schematics of the daughter-daughter interface angle (left) and box plot of the daughter- daughter interface angle (median ± interquartile range, right) at 80% of apical ring contraction in ctrl, *cysts*, *rho1 or rok^RNAi^* cells neighbouring a dividing cell. Note that for the *cysts* quantification, both the dividing and the neighbouring cells can be *cysts* mutant cells. (I) Images of Cysts:GFP (left and right) and MyoII:3xmKate2 (left) in a mosaic tissue with ctrl and *Ecad^RNAi^* interphasic cells. *Ecad^RNAi^* cells are marked by the expression of CAAX:tBFP (not shown), and the yellow dashed line marks the boundary between ctrl and *Ecad^RNAi^* cells. (J) Time-lapse images of Ecad:GFP and Cysts:mKate2 in the absence (left) and the presence of LARIAT (right). Time is set to 0 at the beginning of imaging or photoactivation. Yellow arrowheads: Cysts:mKate2 accumulation within Ecad:GFP depletion regions. The brightness- contrast for the Cysts:mKate2 signal was differentially set between ctrl and LARIAT. The increase in Cysts:mKate2 signal in both conditions is due to the use of the bleach correction function in Fiji on the low Cysts:mKate2 signal. (K) Box plot of the Spearman correlation coefficients between Ecad:GFP and Cysts:mKate2 signals in junctional regions without local Ecad:GFP depletion (full circles) or in junctional regions encompassing an Ecad depletion (open circles) upon LARIAT optogenetic activation. (L) Time-lapse images of MyoII:3xmKate2 and Cysts:GFP in the context of a ctrl dividing cell surrounded by ctrl and *rok^RNAi^* neighbouring cells. *rok^RNAi^* cells are marked by the expression of CAAX:tBFP (not shown), and the yellow dashed line marks the boundary between ctrl and *rok^RNAi^* cells. Yellow arrowheads: Cysts:GFP accumulation in the ctrl neighbouring cells. Yellow open arrowheads: reduced Cysts:GFP accumulation in the *rok^RNAi^* neighbours. Inset: close-ups on Cysts:GFP signal at the rim of the cytokinetic ring. Ratios of Cysts:GFP levels between the *rok^RNAi^* and ctrl control neighbours have a median value of 0.44 and an interquartile range from 0.2 to 0.7 (n=26). Scale bars: 5µm (A-I), 1µm (J and insets in D, I). Mann-Whitney tests: *: *p*<0.05, **: *p*<0.01; ****: *p*<0.0001. See also Figure S4, Movies S3 and S4. For details on quantifications, see STAR Methods.

We first analysed whether Cysts accumulation at the rim of the cytokinetic ring occurred in the neighbouring cells. By analysing dividing cells devoid of Cysts:GFP and neighbouring Cysts:GFP expressing cells, we observed that Cysts:GFP accumulates near the neighbour ingressing cell membrane during the cytokinesis of an adjacent dividing cell (Fig. 2B and Movie S3B). In addition, and as observed for MyoII (Pinheiro et al., 2017) (Fig. 2C), high temporal resolution time-lapse imaging revealed that Cysts:GFP speckles flowed within the ingressing membrane during furrowing (Fig. 2C). Based on these findings, we concluded that Cysts accumulated in cells neighbouring a cell undergoing cytokinesis. We then investigated whether Cysts activates the Rho1 GTPase in the neighbours during cytokinesis. We found that the Rho activity reporter (Ani-RBD:GFP, (Munjal et al., 2015)) is enriched in the neighbouring cells within the ingressing membrane region during cytokinesis, and that this enrichment required Cysts activity in the neighbouring cells (Fig. 2D and Movie S3C). Having found that Cysts promotes Rho1 activity in neighbouring cells at the rim of the cytokinesis ring, we then explored whether Cysts localization is a response to the force produced by the constriction of the cytokinetic ring. We and others have shown that cytokinetic contractile forces are reduced in dividing cells mutant for the *Drosophila* septin *peanut* (*pnut*, (Founounou et al., 2013; Guillot and Lecuit, 2013; Pinheiro et al., 2017). In agreement with the hypothesis that Cysts recruitment is dependent on cytokinetic forces, its accumulation was decreased in cells neighbouring *pnut^RNAi^* dividing cells (Fig. 2E). If Cysts contributes to the response to contractile forces, it should regulate MyoII accumulation in the neighbouring cells, and daughter cell membrane juxtaposition. Indeed, in *cysts* mutant neighbours, there was a decrease in MyoII:3xmKate2 accumulation (Fig. 2F, G and Movie S4)., Neither *cysts* nor *rho1* neighbouring cells abrogated MyoII accumulation to the same extent of *rok* neighbouring cells (Fig. 2G). This concurs with our previous finding that the mechanosensing response is partially regulated by Rho activity (Pinheiro et al, 2017). Last, and in agreement with the reduction in MyoII accumulation (Fig. 2F and 2G), loss of Cysts modulated the juxtaposition of the daughter cells membranes, as manifested by the increase in the angle formed by the ingressing membranes (Fig. 2G, 2H and Movie S4). Together, these observations indicate that the RhoGEF Cysts participates in mechanosensing during cell division to enhance MyoII accumulation and the juxtaposition of the daughter cells membranes.

In adherent cells in culture, the Cysts vertebrate homologue p114RhoGEF, is recruited to sites of Ecad accumulation upon chemical activation of MyoII, or upon ectopic and global stretching of the epithelial monolayer (Acharya et al., 2018). Yet, Ecad is locally decreased in response to endogenous cytokinetic mechanical forces in *Drosophila* (Pinheiro et al., 2017), suggesting the existence of an additional mode of Cysts-dependent mechanosensing. To test whether endogenous cytokinetic force promotes Cysts accumulation by a local decrease of Ecad, we took advantage of two complementary approaches in interphasic cells, to mirror the interphasic status of the neighbouring cells. First, reduction of Ecad levels using *Ecad^RNAi^* mediated knockdown led to an enrichment of Cysts:GFP at cell membranes (Fig. 2I). Interestingly, high-temporal resolution time-lapse imaging showed that Ecad decrease resulted in the formation of both Cysts and MyoII flows towards the ectopic medial MyoII accumulation caused by *Ecad^RNAi^* mediated knockdown (Fig. S4B and S4C). Second, we used the LARIAT system, an optogenetic approach to promote the clustering of GFP tagged proteins (Lee et al., 2014; Qin et al., 2017), to generate local depletions of Ecad with high temporal control. By promoting light-induced clustering of Ecad:GFP in a tissue where Ecad:GFP was the only source of Ecad, we generated local junctional Ecad:GFP depletions, and found that Cysts:mKate2 became enriched at Ecad:GFP depletion sites (Fig. 2J and 2K). Together, these results indicate that Ecad decrease can promote Cysts accumulation. Knowing that MyoII flows and accumulation depend on Rok function (Herszterg et al., 2013; Pinheiro et al., 2017), and having observed that Cysts flows with MyoII within the ingressing membranes (Fig 2C), we also hypothesized that Cysts local accumulation could be reinforced by Rok. We therefore analysed Cysts:GFP accumulation in *rok^RNAi^* neighbouring cells, which are characterized by the lack of MyoII flows and accumulation (Herszterg et al., 2013; Pinheiro et al., 2017). In the absence of Rok function, Cysts accumulation was reduced in the neighbouring cells (Fig 2L). We therefore propose that Ecad dilution in response to cytokinesis forces promotes Cysts recruitment, which is reinforced by Rok activity.

In summary, we propose that the RhoGEF Cysts accumulates at the rim of the cytokinetic ring in response to a local decrease of Ecad promoted by mechanical force. In turn, Cysts is necessary to activate Rho1 and to enhance both MyoII accumulation and membranes juxtaposition prior to *de novo* junction formation; thus, establishing a role for Cysts in the response to endogenous mechanical forces during epithelial cell division.

### RhoGEF4 controls *de novo* AJ length upon cell division

Upon midbody formation, daughter cell membrane juxtaposition is followed by the *de novo* formation of an AJ between the two daughter cells, a feature conserved in multiple epithelial tissues including the notum and the FE (Herszterg et al., 2014; Ragkousi and Gibson, 2014). The length and the topology of *de novo* junctions is regulated in the dividing cell by Rac, which promotes Arp2/3-dependent F-Actin polymerization around the midbody and at the daughter cells interface (Herszterg et al., 2013; Morais-De-Sá and Sunkel, 2013). In particular, F-Actin polymerization propels the withdrawal of the neighbouring cell membranes inserted between the daughter cells; thus enabling the formation of daughter cell junctions (Herszterg et al., 2013). Yet, the mechanisms of Rac activation during epithelial cytokinesis are unknown.

To investigate how *de novo* junctions are formed upon division, we decided to focus on RhoGEF4, since it became enriched at the daughter cells interface and midbody (Fig. 3A, Fig. S5C and Movie S5A), mirroring the accumulation of F-Actin observed in both the notum and the FE (Herszterg et al., 2013; Morais-De-Sá and Sunkel, 2013). The human FGD3/FGD4 orthologues of RhoGEF4 remain poorly characterized despite their putative implications in cancer or Charcot-Marie tooth disease (Delague et al., 2007; Renda et al., 2019). Using the *rhogef4* null allele generated by CRISPR/Cas9 mediated homologous recombination, we found that the initial junctions formed between *rhogef4* daughter cells were shorter in both tissues (Fig. 3B, 3C, Fig. S5D, S5E and Movie S5B). These data prompted us to explore the role of RhoGEF4 during cytokinesis in more detail, and to investigate which RhoGTPase it regulates to control the length of *de novo* junctions formed upon division.

**Figure 3:**
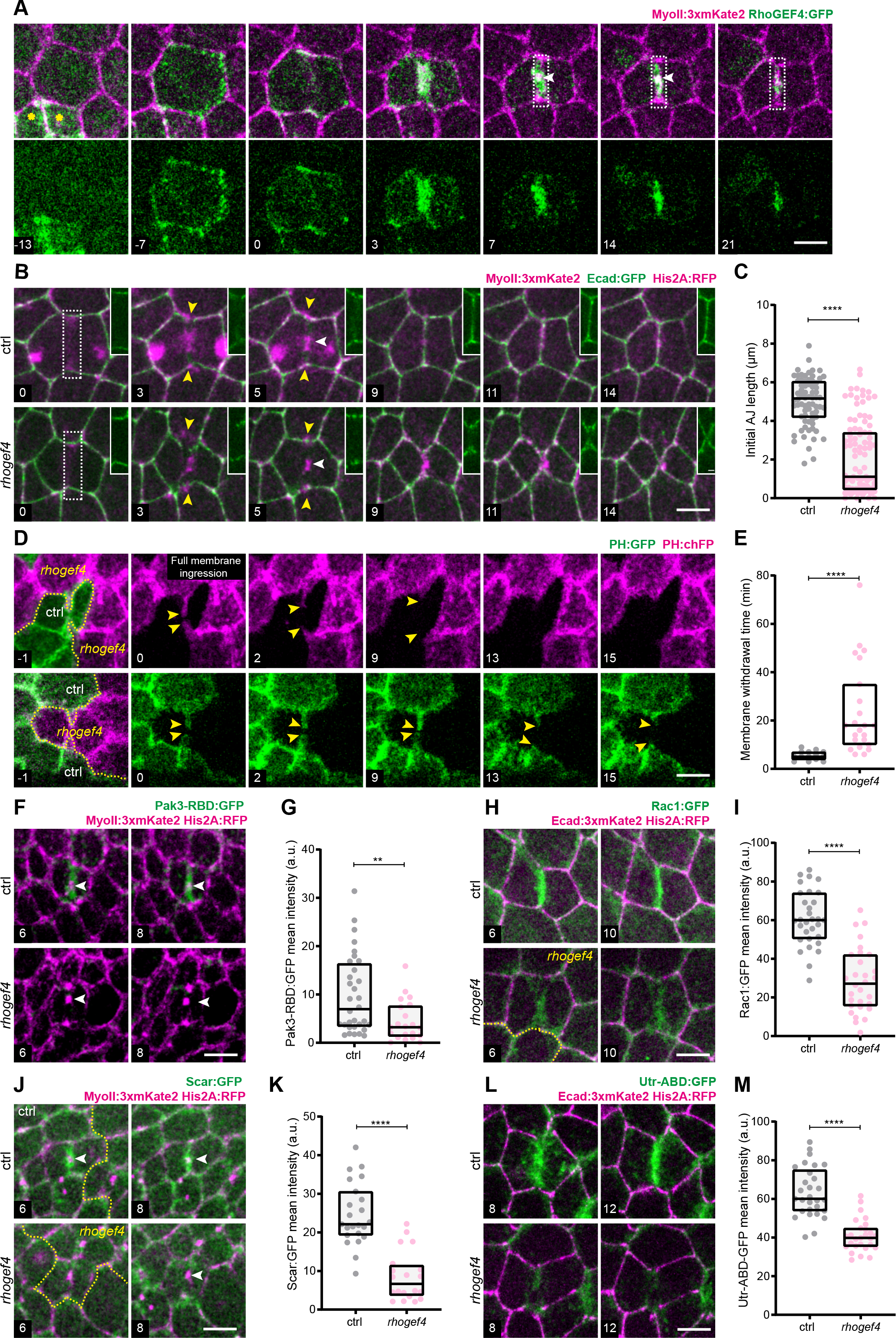
RhoGEF4 regulates *de novo* cell junction length upon cytokinesis. All confocal images are apical top views at the level of the AJ in the notum. Unless otherwise specified, time (min, indicated in the lower left image corner) is set to 0 at the onset of cytokinesis marked by the initial deformation of AJ by the constriction of the cytokinetic ring. In B, F, J and L, ctrl and *rhogef4* cells are respectively marked by the presence and absence of His2A:RFP signal. Since the nuclei can be located basal to the confocal section shown, the His2A:RFP signal is not visible in all panels or ctrl cells. (A) Time-lapse images of RhoGEF4:GFP (top and bottom) and MyoII:3xmKate2 (top) from interphase to late cytokinesis. White arrowheads: midbody. White dashed box: daughter cell interface. The yellow asterisks indicate two neighbouring cells in late cytokinesis. (B) Time-lapse images of Ecad:GFP, MyoII:3xmKate2 and His2A:RFP in ctrl and *rhogef4* dividing cells. *rhogef4* cells are marked by the absence of His2A:RFP expression. Inset: close-up on the Ecad:GFP signal along the daughter cell interface formed during cytokinesis, in the region marked by the white dashed box. White arrowheads: midbody. Yellow arrowheads: MyoII accumulation in the neighbouring cells and prospective position of *de novo* daughter-daughter cell junction. (C) Box plot of *de novo* AJ length (median ± interquartile range) in ctrl and *rhogef4* cells. (D) Time-lapse images of PH:GFP and PH:chFP in a ctrl dividing cell (marked by PH:GFP) surrounded by *rhogef4* cells (marked by PH:chFP, top), and in a *rhogef4* dividing cell (marked by PH:chFP) surrounded by ctrl neighbouring cells (marked by PH:GFP, bottom). Time (min) is set to 0 at the time of full neighbouring membrane ingression. The dividing cell membranes are only shown in the first panel. *rhogef4* cells are marked by the absence of PH:GFP expression, and the yellow dashed line outlines the boundary between *rhogef4* and ctrl cells. Yellow arrowheads: position of the tip of the neighbouring membranes inserted between the two daughter cells. (E) Box plot of the duration of neighbouring cell membrane withdrawal (median ± interquartile range) from ctrl or *rhogef4* daughter cell apical interfaces. (F) Time-lapse images of MyoII:3xmKate2 and Pak3-RBD:GFP for ctrl and *rhogef4* dividing cells during cytokinesis. White arrowheads: midbody. (G) Box plot of Pak3-RBD:GFP intensity (median ± interquartile range) at the apical daughter cell interface in ctrl and *rhogef4* cells. (H) Time-lapse images of Ecad:3xmKate2 and Rac1:GFP in ctrl and *rhogef4* dividing cells during cytokinesis. The yellow dashed line outlines the boundary between *rhogef4* and ctrl cells. (I) Box plot of Rac1:GFP intensity (median ± interquartile range) at the apical daughter cell interface in ctrl or *rhogef4* daughter cells during late cytokinesis. (J) Time-lapse images of Scar:GFP and MyoII:3xmKate2 in ctrl and *rhogef4* dividing cells during cytokinesis. White arrowheads: midbody. The yellow dashed line outlines the boundary between *rhogef4* and ctrl cells. (K) Box plot of Scar:GFP intensity (median ± interquartile range) at the apical daughter cell interface in ctrl or *rhogef4* daughter cells. (L) Time-lapse images of Ecad:3xmKate2 and Utr-ABD:GFP in ctrl and *rhogef4* dividing cells during cytokinesis. (M) Box plot of Utr-ABD:GFP intensity (median ± interquartile range) at the apical daughter cell interface in ctrl or *rhogef4* daughter cells. Scale bars: 5µm (A, B, D, F, H, J, L), 1µm (inset in B). Mann-Whitney test: ****: *p*<0.0001. **: p<0.01. See also Figure S5 and Movie S5. For details on quantifications, see STAR Methods.

As RhoGEF4 localizes to different structures during cell division (Fig. 3A, Fig. S5A- S5C and Movie S5A), we first tested whether RhoGEF4 specifically acts at the newly forming junction. During cytokinesis, RhoGEF4 localizes at the cytokinetic ring and thus, near the tip of the ingressing daughter cell membranes. We found nevertheless that the loss of RhoGEF4 function did not affect the juxtaposing of the daughter cell membranes during cytokinetic ring contraction (Fig. S5F). Since RhoGEF4 localized at the cytokinetic ring and at the reorganizing nuclear envelope during cytokinesis, we then tested whether RhoGEF4 contributes to ring contraction and nuclear envelope dynamics. There were no defects observed in the ring contraction rate of *rhogef4* dividing cells (Fig. S5G). Moreover, the dynamics of both Lamin:TagRFP and nls:GFP relocalization to the nucleus were similar in control and *rhogef4* telophase cells (Fig. S5H and S3I). Together, these data indicate that RhoGEF4 does not regulate daughter membrane juxtaposition, ring contraction and nuclear envelope reformation, prompting us to explore how it specifically contributes to *de novo* AJ length regulation during late cytokinesis.

Towards this goal, we first analysed Ecad dynamics during *de novo* junction formation. We found that the loss of RhoGEF4 function often caused a delay in the initial Ecad enrichment at the presumptive daughter cell interface in the notum (Fig. 3B, FigS5J). Yet, and as previously found for the loss of function of Arp3 in the notum (Herszterg et al., 2013), the occurrence of this delay did not correlate with a reduced junction length (Fig. S5J). In addition, the timing of Ecad accumulation was unaffected in *rhogef4* cells in the FE (Fig. S5K). Together, these findings indicate that RhoGEF4 regulates AJ length independently of the accumulation dynamics of Ecad at the daughter interface. We therefore hypothesized that RhoGEF4 regulates

AJ length by controlling the withdrawal of neighbouring cell membranes before junction formation. The dynamics of the daughter and neighbour cell membranes can be tracked by labelling the dividing and neighbour cell membranes with PH:GFP and PH:chFP, respectively. Using this approach, we found that the membrane withdrawal of cells neighbouring *rhogef4* dividing cells was significantly delayed relative to control dividing cells (Fig. 3D and 3E). We therefore concluded that in the dividing cell RhoGEF4 promotes the withdrawal of the neighbour membranes thereby controlling *de novo* AJ length.

RhoGEF4 can act as a GEF for Rac or Rho *in vitro* (Nahm et al., 2006). We therefore tested whether RhoGEF4 regulates Rac or Rho1 activities in the control of neighbouring membrane withdrawal. Towards this end, we compared the dynamics of the Rho activity reporter Ani-RBD:GFP, (Munjal et al., 2015), and of a Rac activity reporter (Pak3-RBD:GFP, (Abreu-Blanco et al., 2014)) during cytokinesis. In control dividing cells, Ani-RBD:GFP was strongly enriched at the contractile ring and the midbody (Fig. S5L), whereas Pak3-RBD:GFP was present at the daughter cell interface before *de novo* junction formation (Fig. 3F). While Ani-RBD:GFP localization was unaffected in *rhogef4* cells (Fig. S5L and S5M), Pak1- RBD:GFP signal was reduced at the daughter cell interface in *rhogef4* cells (Fig. 3F and 3G). Further confirming the role of RhoGEF4 in controlling Rac function, we found that (i) the loss of RhoGEF4 function led to a decrease of Rac1:GFP at the daughter cell interface (Fig. 3H and 3I, Movie S5C); and (ii) Scar, a downstream effector of Rac and an Arp2/3 activator (Georgiou and Baum, 2010), was also decreased at the daughter cell interface of *rhogef4* cells (Fig. 3J and 3K). Finally, as observed upon reduction of Rac function (Herszterg et al., 2013), the Utrophin- ABD:GFP F-Actin probe was strongly reduced at the daughter cell interface in *rhogef4* dividing cells (Fig. 3L,M, Movie S5D). Together, these data indicate that RhoGEF4 acts in the dividing cell to regulate *de novo* AJ length by controlling via Rac the F-actin accumulation at the daughter cell interface to induce neighbouring cell membrane withdrawal.

### RhoGEF4 and Cysts cooperate in defining the initial *de novo* junction topology

Upon cytokinesis, two distinct topological AJ arrangements have been reported in different epithelia: a daughter-daughter (d-d) AJ or neighbour-neighbour (n-n) AJ (Aigouy et al., 2010; Firmino et al., 2016; Gibson et al., 2006; Guillot and Lecuit, 2013; Herszterg et al., 2014; Higashi et al., 2016; Lau et al., 2015; Morais-De-Sá and Sunkel, 2013; Ragkousi and Gibson, 2014; Uroz et al., 2018). Having found that RhoGEF4 is a key regulator of the length of AJ between daughter cells, we analysed whether the loss of RhoGEF4 modulates the topology of cell arrangements upon cytokinesis. Whereas in wt cells, a d-d AJ is always directly formed upon cell division (n=120), we found that in a fraction of *rhogef4* cells in the notum (11%, n=120), an AJ is transiently formed between the neighbouring cells instead of the two daughter cells (Fig. 4A, Fig. S6A, S6D and Movie S6). Importantly, the formation of n-n AJ did not correlate with the timing of Ecad enrichment at the initial interface (Fig. S6A) and was rather associated with the formation of an initial very short d-d contact (i.e., less than 1 µm). We also found that the impact of RhoGEF4 loss of function on the length of the d-d junction was stronger in the notum than in the FE, where the very short d-d junctions that could promote topological defects were unfrequently observed (2 cases out of 19 divisions). While this could be due to compensatory mechanisms associated with the function of other RhoGEF localized at the d-d interface in the FE, we recalled that in contrast to the FE, the notum undergoes major morphological changes during proliferation (Chen et al., 2019; Duhart et al., 2017; Guirao et al., 2015). We thus hypothesized that in tissues undergoing major morphogenesis movements, the loss of RhoGEF4 function could lead to the formation of very short junctions, and thus transient n-n junctions. We therefore analysed the impact of RhoGEF4 loss of function in the pupal histoblast and wing, two epithelial tissues characterized by proliferation and large morphogenetic movements (Aigouy et al., 2010; Bischoff and Cseresnyes, 2009; Davis et al., 2022; Guirao et al., 2015; Ninov et al., 2009; Sugimura and Ishihara, 2013). Strikingly, in both tissues, the loss of RhoGEF4 function led to very short AJ formation and topological defects during cell division (Fig. S6B-D). So far, the topology of AJ formed upon cell division has been proposed to regulate tissue dynamics and planar cell polarity (Aigouy et al., 2010; Firmino et al., 2016; Lau et al., 2015); our results support the notion that the overall dynamics of epithelial tissues can also contribute to the initial topology of the junctions formed upon cytokinesis.

**Figure 4:**
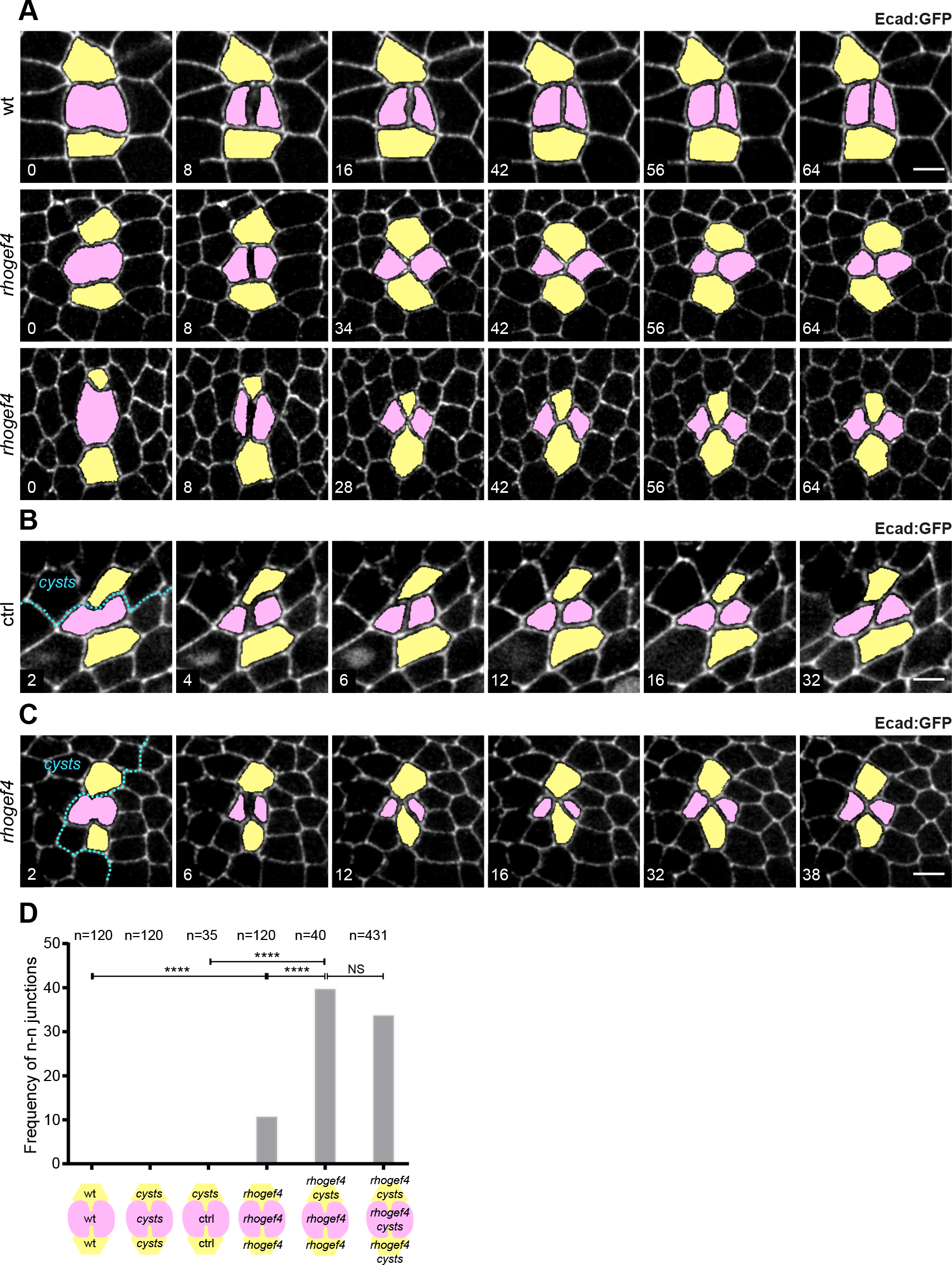
RhoGEF4 and Cysts synergistically control junction topology upon cell division. All images are apical top views of the notum epithelium at the level of the AJ labelled by Ecad:GFP. The dividing and daughter cells are coloured in pink while the neighbouring cells are in yellow. Unless otherwise indicated, time (min, indicated in the lower left image corner) is set to 0 at the onset of cytokinesis marked by the initial deformation of AJ by the constriction of the cytokinetic ring. In B and C, ctrl and *cysts* cells are respectively marked by the presence and absence of nls:GFP signal. Since the nuclei can be located basal to the confocal section shown, the nls:GFP signal is not visible in all panels or ctrl cells. (A) Time-lapse images of Ecad:GFP from cytokinesis onwards in wt (top) or *rhogef4* (middle and bottom) pupae. Two examples are shown for *rhogef4* tissues: in the middle panels, a d-d junction is formed upon cell division (from t = 34 min onwards), whereas in the bottom ones a n-n junction is formed (from t = 30 min onwards). (B) Time-lapse images of Ecad:GFP from cytokinesis onwards of a control (ctrl) dividing cell neighbouring *cysts* or ctrl cells. *cysts* cells are marked by the absence of nls:GFP signal, and the cyan dashed line outlines the boundary between *cysts* and ctrl cells. (C) Time-lapse images of Ecad:GFP from cytokinesis onwards of a *rhogef4* dividing cell neighbouring *rhogef4, cysts* or *rhogef4* cells. *cysts* cells are marked by the absence of nls:GFP signal, and the cyan dashed line outlines the boundary between *cysts,rhogef4* and *rhogef4* cells. (D) Graph of the frequency of n-n junctions formed upon cell division in wt, *cysts*, ctrl, *rhogef4*, or double *cysts*, *rhogef4* mutant dividing cells neighbouring wt, *cysts*, ctrl, *rhogef4* or double *rhogef4*, *cysts* mutant cells, respectively, as indicated in the schematics below the graph. Scale bars: 5µm. Zscore proportion test: NS: non significant, ****:*p* ≤0.0001. See also Figure S6 and Movie S6. For details on quantifications, see STAR Methods.

Having identified two RhoGEF respectively regulating membranes juxtaposition during ring contraction, and membrane withdrawal upon midbody formation, we could then investigate, for the first time, whether these two processes cooperate to regulate the topology of the newly formed AJ. Towards this goal, we analysed whether in *rhogef4* tissues, the proportion of topological defects upon cytokinesis increases when *rhogef4* dividing cells are neighboured by a cell also mutant for *cysts*. In these conditions, the proportion of n-n AJ increased by 3.6-fold relative to *rhogef4* dividing cells neighbouring cells devoted of *cysts* mutation (Fig. 4B-D). Since control dividing cells neighboured by a *cysts* cell never formed n-n AJ (0%, n=35, Fig. 4B and 4D), our results revealed a synergistic role of RhoGEF4 and Cysts in the control of AJ topology during cytokinesis. Together, we conclude that daughter cell membrane juxtaposition facilitated by Cysts via Rho, and neighbouring membrane withdrawal mediated by RhoGEF4 via Rac coregulate *de novo* junction formation and topology during epithelial cell division *in vivo*.

## DISCUSSION

By modulating the activity of small RhoGTPases, RhoGEF and RhoGAP control cytoskeleton organization and dynamics in a broad range of processes such as cell morphogenesis, division and migration in all eukaryotes (Arnold et al., 2017; Buckley and St Johnston, 2022; Denk-Lobnig and Martin, 2019; Heasman and Ridley, 2008; Iden and Collard, 2008; Jaffe and Hall, 2005). The functions of RhoGTPases have been extensively and systematically investigated in individual cells. However, multicellularity entails a complex interplay between the cytoskeleton and cell-cell junctions to regulate cell and tissue architecture and dynamics (Buckley and St Johnston, 2022; Fernandez-Gonzalez and Peifer, 2022); thus highlighting the relevance of characterizing RhoGEF/GAP function in tissues. Cell division has emerged as a multicellular process, since it entails the deformation of the neighbouring cells, the remodelling of the dividing and neighbour cell junctions, as well as *de novo* junction formation (Herszterg et al., 2014; Ragkousi and Gibson, 2014). To better understand how RhoGEF and RhoGAP control the cytoskeleton and cell junction dynamics in proliferative epithelial tissues, we have generated a family-wide *Drosophila* transgenic library of fluorescently tagged RhoGEF/GAP, and systematically determined their distributions during interphase and cell division in two epithelial tissues. Our screen revealed multiple uncharacterized RhoGEF/GAP interphasic localizations as well as a complex choreography of RhoGEF/GAP during cell division. Building on our screen, we have better defined the processes of mechanosensing and junction formation, by delineating how the activities of specific RhoGTPases are controlled during epithelial cytokinesis and *de novo* junction formation. We have therefore uncovered the first mechanisms of RhoGTPases activation underlying the multicellularity of epithelial cell division.

It is now well established that cytokinesis and more generally cell division entail a mechanosensing response in the neighbours (Higashi et al., 2016; Monster et al., 2021; Pinheiro et al., 2017). Despite such conserved mechanical responses and the proposed roles of RhoGTPases in mechanosensing (Pinheiro and Bellaïche, 2018), the nature of the RhoGEF associated with the response to cytokinesis ring contraction has remained unknown. Our work establishes that the RhoGEF Cysts modulates the response to endogenous mechanical forces generated by ring contraction in the neighbouring cells, thereby enhancing daughter cell membranes juxtaposition. Interestingly, whereas previous studies in cell culture showed that the Cysts orthologue p114RhoGEF is recruited at sites of Ecad accumulation (Acharya et al., 2018), we uncovered an additional mechanism of Cysts localization promoted by local Ecad decrease due to mechanical forces. To further understand how cells respond to mechanical forces associated with a local Ecad decrease, it will be relevant to explore how Cysts is recruited to sites of lower Ecad, and how Rok-dependent contractility feedbacks on Cysts dynamics. Interestingly, at least two distinct Ecad dynamics have been observed during epithelial cytokinesis in vertebrates (Firmino et al., 2016; Higashi et al., 2016). In the avian embryo at stage X, cell division is concomitant to a local depletion of basal Ecad and an accumulation of MyoII and F-Actin (Firmino et al., 2016), whereas in Xenopus, ring contraction triggers a well-known force sensing response associated with the accumulation of Ecad and Vinculin (Higashi et al., 2016). Building on our findings on Cysts and the ones on p114RhoGEF *in vitro* (Acharya et al., 2018), one could hypothesize that Cysts/p114RhoGEF represents a common platform for the response to mechanical force during cytokinesis; thus calling for the investigation of the function of RhoGTPases in cells neighbouring the dividing cells in vertebrates. Last, Cysts is involved in AJ integrity, cell-cell rearrangements and cell ingression in interphasic cells in the early *Drosophila* embryo (Garcia De Las Bayonas et al., 2019; Silver et al., 2019; Simões et al., 2022); processes entailing the production of mechanical forces and the remodelling of the actomyosin cytoskeleton (Fernandez-Gonzalez and Peifer, 2022; Lecuit and Yap, 2015; Pinheiro and Bellaïche, 2018). One could envision that cytoskeleton and junction dynamics might also depend on the Cysts mechanosensing activity in these developmental processes.

*De novo* junction formation is inherent to the proliferation of epithelia. *In vitro* experiments have put forward a fundamental role for Rac in modulating the dynamics of junction formation upon cell-cell contact formation (Chu et al., 2004; Verma et al., 2012; Yamada and Nelson, 2007). Previous findings have also underscored a role for Rac activity in cytokinesis regulating both ring constriction and the topology of the daughter-daughter cell interface (Herszterg et al., 2013; Loria et al., 2012). Here, we uncovered that RhoGEF4 promotes the localization of active Rac at the daughter-daughter cell interface to ensure the withdrawal of the neighbouring cell membranes. Thereby, RhoGEF4 is a regulator of Rac function in the control of *de novo* junction length as well as of the dynamics of the cell-cell arrangements upon cytokinesis. By comparing the role of RhoGEF4 in different tissues, our work also suggests that the initial topology of the junction formed upon cytokinesis can be modulated by the overall dynamics or mechanical properties of epithelial tissues. Last, by combining loss of Cysts and RhoGEF4 function, we establish that the topology of *de novo* junctions formed upon cell division is synergistically regulated both by membranes juxtaposition during ring constriction, and by membranes withdrawal upon midbody formation. Altogether, our analyses of Cysts and RhoGEF4 functions provide a far better understanding of the mechanisms of Rho and Rac regulation during epithelial cell division, and they illustrate how their distinct activations in the dividing and neighbouring cells couple cytokinesis and junction formation.

Our findings on Cysts and RhoGEF4 also highlight the relevance of screening fluorescently tagged libraries to pinpoint key regulators of small GTPases necessary for cytoskeleton and junctional regulations *in vivo*. In particular, we foresee that family wide characterization of RhoGEF/GAP localization and dynamics will be instrumental to better understand numerous aspects of the conserved process of cell division in invertebrates and vertebrates. So far, studies on cell division have mainly focused on the roles of two RhoGTPase regulators, ECT2 (*Drosophila* Pbl) and RACGAP1 (*Drosophila* Tum) (Glotzer, 2017; Mishima et al., 2002; Su et al., 2011; Yüce et al., 2005; Zhao and Fang, 2005). Our systematic analysis now uncovered a very diverse set of RhoGEF/GAP localizations during epithelial cell division. Accordingly, our screen suggested several possible avenues to decipher the function of RhoGEF/GAP in mitotic rounding, polar relaxation and asymmetric furrowing as well as in midbody dynamics. We also observed that several RhoGEF/GAP display similar localizations. These findings will be instrumental to design double or triple loss of function experiments to explore RhoGEF/GAP functions, and to complement loss of function screens that would overlook specific RhoGEF/GAP functions due to their functional redundancy. In addition, our analysis in two distinct tissues underscores a set of RhoGEF/GAP with different localization and dynamics. This illustrates the relevance of the *in vivo* exploration under the control of the endogenous promoters to explore how mitosis and cytoskeleton dynamics are differentially modulated to regulate tissue development or function. More generally, RhoGTPases are central regulators of cell and tissue dynamics in metazoans (Arnold et al., 2017; Heasman and Ridley, 2008; Jaffe and Hall, 2005); we therefore expect that the library will be a key resource to dissect RhoGTPases spatiotemporal regulation and function in a variety of developmental, homeostatic and repair contexts in epithelia as well as in stem cells, migrating cells and neurons.

## Supporting information

Key Ressource Table

## AUTHOR CONTRIBUTIONS

Experiments: F.d.P., M.O., J.H., I.C., J.L-G., Z.W.; Data Analysis: F.d.P., M.O., J. H, I.C., J.L-G.; Methodology: C.M., I.G., S.P.; Resources: I.G., S.P, C.M., J. H, M.O., F.d.P, A.L.Visualization: F.d.P., M.O., E.M.S.; Writing – original draft: F.d.P., I.C. Y.B., E.M.S.; Writing – review & editing: F.d.P., Y.B., M.O., I.C., E.M.S., S.P., J. H; Supervision: Y.B and E.M.S; Project administration: Y.B. and E.M.S; Funding acquisition: Y.B., F.d.P. and E.M.S.

## ACKNOWLEDGEMENTS

We thank T. Lecuit, M. Suzanne, F. Pichaud, the Bloomington Stock Centre, Vienna Drosophila Resource Center, Transgenic RNAi Project at Harvard Medical School, Kyoto Stock Center for fly lines; L. Alpar, M. Balakireva, F. Bosveld for critical comments on the manuscript; F. Macaluso for help with experiments characterizing RhoGEF4 function; A.F. Santos and A. Barros-Carvalho for help with the production of RhoGEF lines; the PICT-IBiSA@BDD imaging facility of Institut Curie (member of the French National Research Infrastructure France-BioImaging, ANR-10-INBS-04); the i3S Scientific Platform ALM, member of the national infrastructure PPBI - Portuguese Platform of Bioimaging (PPBI-POCI-01-0145- FEDER-022122). ERC Advanced (TiMorph, 340784), ARC (SL220130607097), ANR (TiMecaDiv 20CE13000801), CANCERO-INCA (PLBIO2020/BELLAICHE), ANR Labex DEEP (11-LBX-0044, PSL ANR-10-IDEX-0001-02) grants for funding. FCT—Fundação para a Ciência e a Tecnologia, I.P., under the projects PTDC/BIA-CEL/1511/2021 and PTDC/BEX-BCM/0432/2014. F.d.P. was supported by a FRM post-doctoral fellowship (SPF20170938661). M.O. was supported by a PhD fellowship from FCT and the post-doctoral salary by the Maria de Sousa Award (48/2021).

---

**Figure S1:**
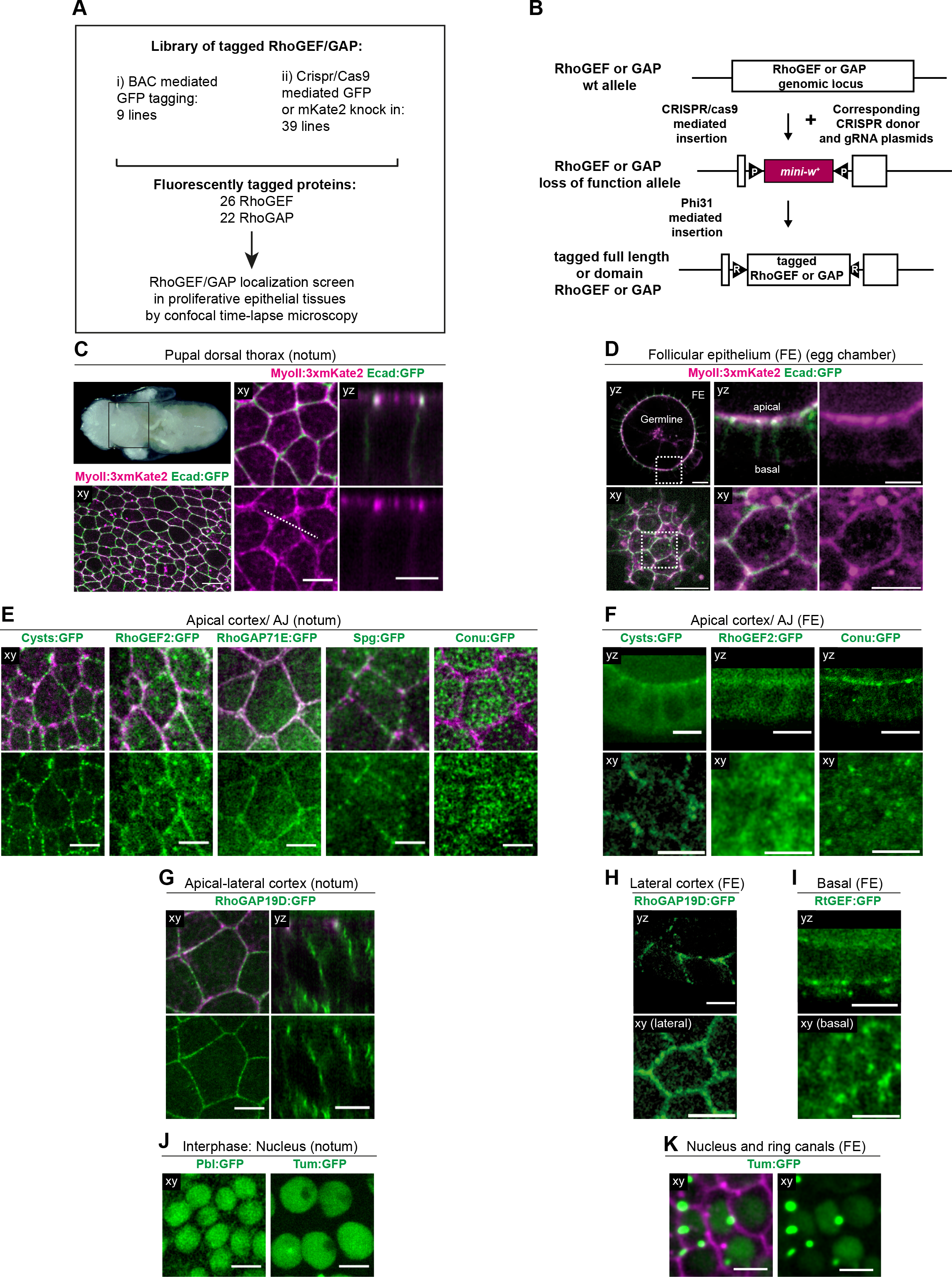
Libraries and tagged RhoGEF/GAP localizations in interphase, related to Figure 1. Unless otherwise indicated, in all images, ‘xy’ indicates apical confocal top views at the level of the AJ in the notum and FE, while ‘yz’ denotes apical-basal confocal sections for the notum, and midsagittal view of the egg chamber. MyoII:3xmKate2 is shown in magenta. See also Fig. S2, Tables S1 and S2.. (A) Overview of the transgenic lines generated to characterize the fluorescently tagged RhoGEF/GAP localizations. (B) Schematics describing the use of the donor and guide plasmid library to generate RhoGEF/GAP loss of function alleles that can be modified to produce tagged or sequence specific alleles. (C) Dorsal view of *Drosophila* pupa at 14 hours after pupa formation (hAPF) (top, left from (Bosveld et al., 2012)). The black box in the top panel indicates the whole notum tissue. xy image of MyoII:3xmKate2 and Ecad:GFP in the notum (bottom, left). xy (middle) and yz (right) images of the MyoII:3xmKate2 (top and bottom) and Ecad:GFP (top) distributions in the notum. Dashed line: position of the yz section. (D) Images of MyoII:3xmKate2 and Ecad:GFP (left and middle) in midsagittal (yz, top) and surface (xy, bottom) views of a stage 4 egg chamber. The middle and right panels correspond to close-ups in the regions outlined in the left panels. (E) xy images of Cysts:GFP, RhoGEF2:GFP, RhoGAP71E:GFP, Spg:GFP, Conu:GFP (top and bottom) and MyoII:3xmKate2 (top) in the notum. (F) yz (top) and xy (bottom) images of Cysts:GFP, RhoGEF2:GFP and Conu:GFP in the FE. (G) xy (left) and yz (right) images of RhoGAP19D:GFP (top and bottom) and MyoII:3xmKate2 (top) in the notum. (H) yz (top) and xy (bottom) images of RhoGAP19D:GFP in the FE. (I) yz (top) and xy (bottom) images of RtGEF:GFP in the FE. (J) xy images at the level of the nucleus of Pbl:GFP and Tum:GFP in the notum. (K) xy images of Tum: GFP (left and right) and Cell membrane marker (right) in the FE. Images correspond to projection of the nuclei and ring canals. Scale bars: 5µm

**Figure S2:**
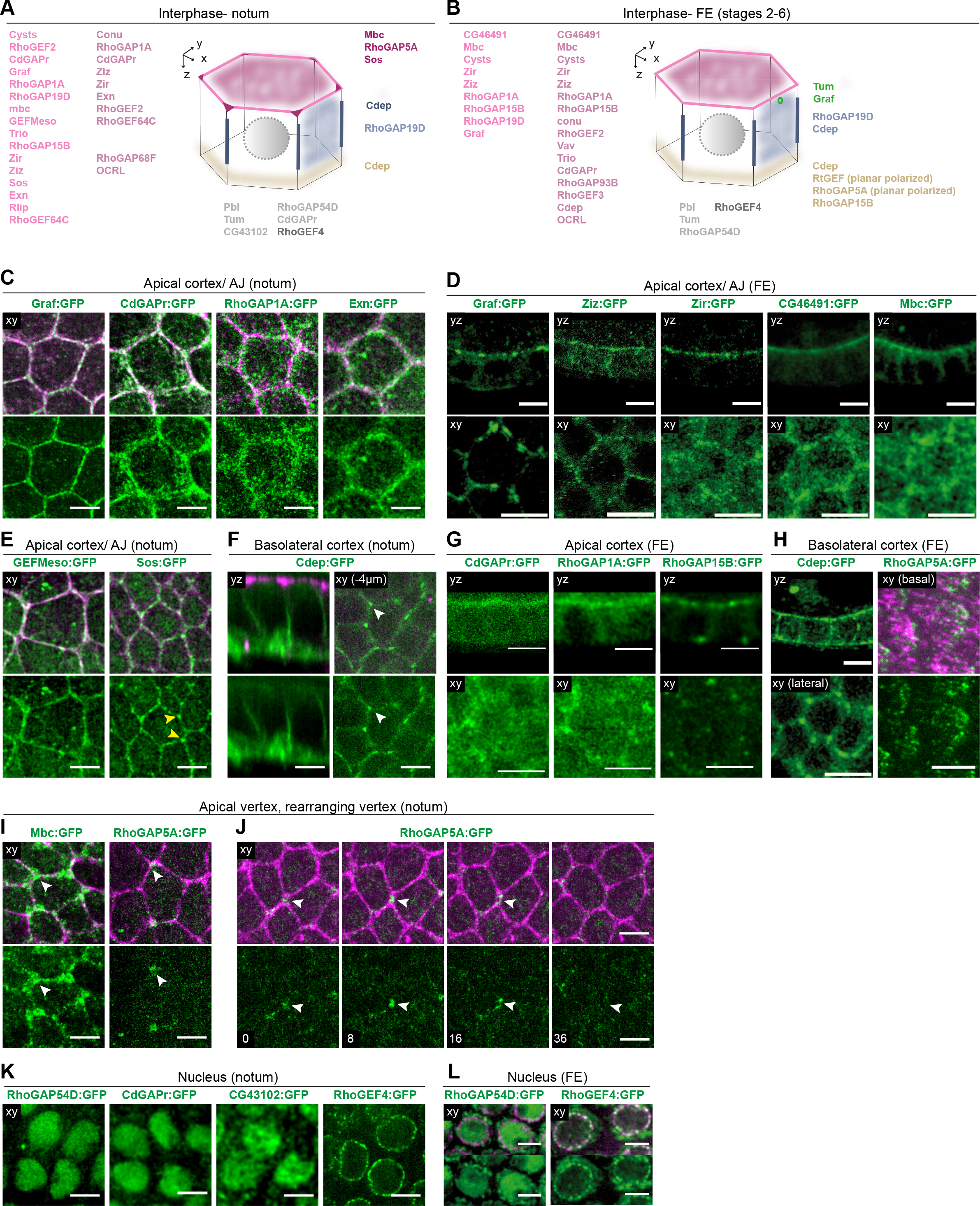
L**o**calizations **of tagged RhoGEF/GAP in interphase, related to** Figure 1. Unless otherwise indicated, for all images, ‘xy’ indicates apical confocal top views at the level of the AJ in the notum and FE, while ‘yz’ denotes apical-basal confocal sections for the notum, and midsagittal views of the FE. MyoII:3xmKate2 is shown in magenta. (**A, B**) Schematics of a notum (A) and a FE (B) epithelial cell with the subcellular localizations of RhoGEF/GAP in interphase. RhoGEF/GAP are color-coded according to their localization. Only the RhoGEF/GAP observed with a sufficient signal to noise ratio in the notum or the FE are shown. (C) xy images of Graf:GFP, CdGAPr:GFP, RhoGAP1A:GFP, Exn:GFP (top and bottom) and MyoII:3xmKate2 (top) in the notum. (D) yz (top) and xy (bottom) images of Graf:GFP, Ziz:GFP, Zir:GFP, CG46491:GFP and Mbc:GFP in the FE. (E) xy images of GEFMeso:GFP, Sos:GFP (top and bottom) and MyoII:3xmKate2 (top) in the notum epithelium. Yellow arrowheads: Sos:GFP accumulations at the apical TCJ. (F) yz (left) and xy (right, confocal section taken 4µm below the AJ) images of Cdep:GFP (top and bottom) and MyoII:3xmKate2 (top) in the notum. White arrowhead: Cdep enrichment at basolateral TCJ. (G) yz (top) and xy (bottom) images of CdGAPr:GFP, RhoGAP1A:GFP and RhoGAP15B:GFP in the FE. (H) yz (left, top) and xy (left, bottom) images of Cdep:GFP; xy images of RhoGAP5A:GFP (right, top and bottom), and MyoII:3xmKate2 (right, top) in the basal cortex of the FE. (I) xy images of Mbc:GFP, RhoGAP5A:GFP (top and bottom) and MyoII:3xmKate2 (top) in the notum. White arrowheads: Mbc:GFP and RhoGAP5A:GFP accumulations at the apical TCJ. (J) xy time-lapse images of RhoGAP5A:GFP (top and bottom) and MyoII:3xmKate2 (top) during the junction elongation phase of a cell-cell rearrangement in the notum. White arrowheads: RhoGAP5A:GFP accumulation. Time is in min. (K) xy images at the level of the nucleus of RhoGAP54D:GFP, CdGAPr:GFP, CG43102:GFP and RhoGEF4:GFP in the notum. (L) xy images at the level of the nucleus of RhoGAP54D:GFP, RhoGEF4:GFP (top and bottom) and Nup107:RFP (top) in the FE. Scale bars: 5µm See also Fig. S1, Table S2..

**Figure S3:**
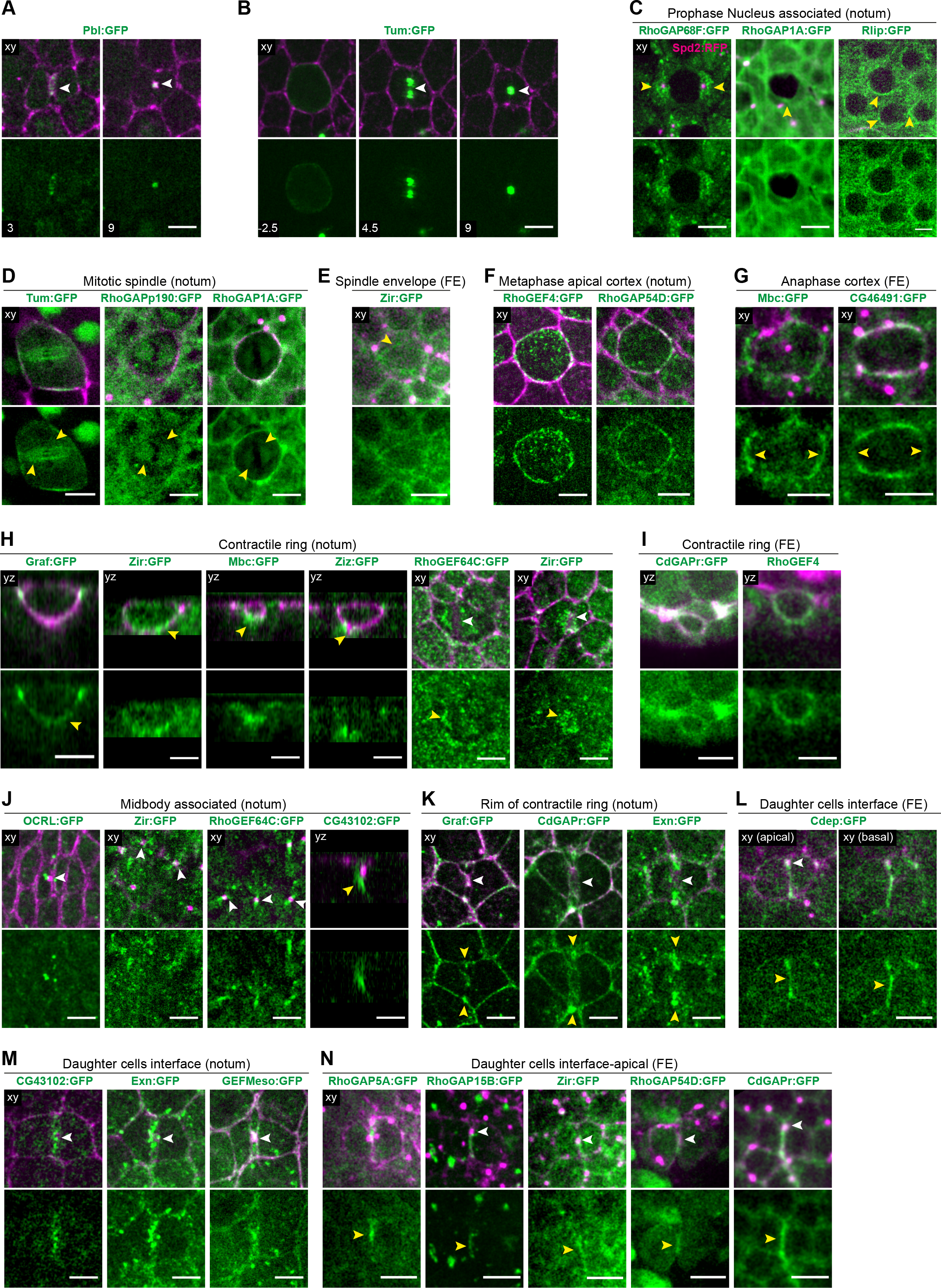
Localizations of additional RhoGEF/GAP during epithelial cell division, related to Figure 1. Unless otherwise indicated, ‘xy’ indicates apical confocal top views at the level of the AJ in the notum and FE, while ‘yz’ denotes apical-basal confocal section at the level of the cytokinetic ring or the midbody for the notum, and midsagittal view of the egg chamber. MyoII:3xmKate2 is shown in magenta. (A) xy time-lapse images of Pbl:GFP (top and bottom) and MyoII:3xmKate2 (top) during cell division in the notum. Time (min, indicated in the lower left image corner) is set to 0 at the onset of cytokinesis marked by the initial deformation of AJ by the constriction of the cytokinetic ring. (B) xy time-lapse images of Tum:GFP (top and bottom) and MyoII:3xmKate2 (top) during cell division in the notum. White arrowheads: Position of the contractile ring and midbody respectively. Time (min, indicated in the lower left image corner) is set to 0 at the onset of cytokinesis marked by the initial deformation of AJ by the constriction of the cytokinetic ring. (C) xy images at level of the nucleus of RhoGAP68F:GFP, RhoGAP1A:GFP, Rlip:GFP (top and bottom), Spd2:RFP (top, left) and MyoII:3xmKate2 (top middle and top right). Spd2:RFP labels centrosomes. Yellow arrowheads: accumulations of RhoGAP68F around the centrosomes, and accumulations of RhoGAP1A:GFP and Rlip:GFP around the nucleus. (D) xy images at the level of the spindle of Tum:GFP, RhoGAPp190:, RhoGAP1A (top and bottom) and Myo:3xmKate2 (top) in mitotic cells in the notum. Yellow arrowheads indicate GFP accumulations at the spindle. (E) xy images at the level of the spindle of Zir:GFP (top and bottom) and Myo:3xmKate2 (top) in mitotic cells in the FE. Yellow arrowheads indicate Zir:GFP enrichment around the spindle. (F) xy images of RhoGEF4:GFP, RhoGAP54D:GFP (top and bottom) and MyoII:3xmKate2 (top) during metaphase in the notum. (G) xy images of Mbc:GFP and CG46491:GFP (top and bottom) and MyoII:3xmKate2 (top) during anaphase in the FE. Yellow arrowheads: Mbc:GFP enrichment or CG46491:GFP depletion at polar regions of the cortex. (H) yz images of Graf:GFP, Zir:GFP, Mbc:GFP and Ziz:GFP, and xy images of RhoGEF64C:GFP, Zir:GFP (top and bottom) and MyoII:3xmKate2 (top) during cytokinetic ring constriction in the notum. White arrowheads: contractile ring. Yellow arrowheads: Graf:GFP and Zir:GFP accumulations at the cytokinetic ring; Mbc:GFP and Ziz:GFP accumulations below the cytokinetic ring; RhoGEF64C:GFP and Zir:GFP accumulations around the apical contractile ring. (I) yz images of CdGAPr:GFP, RhoGEF4 (top and bottom) and MyoII:3xmKate2 (top) during cytokinetic ring constriction in the FE. (J) xy images of OCRL:GFP, Zir:GFP, RhoGEF64C:GFP and yz images of CG43102:GFP (top and bottom), and MyoII:3xmKate2 (top) during late cytokinesis in the notum. White arrowheads: midbody. Yellow arrowheads: CG43102:GFP accumulation basal to the midbody. (K) xy images of Graf:GFP, CdGAPr:GFP, Exn:GFP (top and bottom) and MyoII:3xmKate2 (top) during cytokinesis in the notum. White arrowheads: cytokinetic ring. Yellow arrowheads: Graf:GFP, CdGAPr:GFP and Exn:GFP accumulations at the rim of the cytokinetic ring. (L) Apical (left) and basal (right) xy images of Cdep:GFP (top and bottom) and MyoII:3xmKate2 (top) during daughter junction formation in the FE. White arrowheads: midbody. Yellow arrowheads: Cdep:GFP accumulation along the daughter interface from the apical to the basal level. (M) xy images of CG43102:GFP, Exn:GFP, GEFMeso:GFP (top and bottom) and MyoII:3xmKate2 (top) during late cytokinesis and *de novo* daughter cell junction formation in the notum. White arrowheads: midbody. (N) xy images of RhoGAP5A:GFP, RhoGAP15B:GFP, Zir:GFP, RhoGAP54D:GFP, CdGAPr (top and bottom) and MyoII:3xmKate2 (top) during late cytokinesis in the FE. White arrowheads: midbody. Yellow arrowheads: new interface. Scale bars: 5 µm

**Figure S4:**
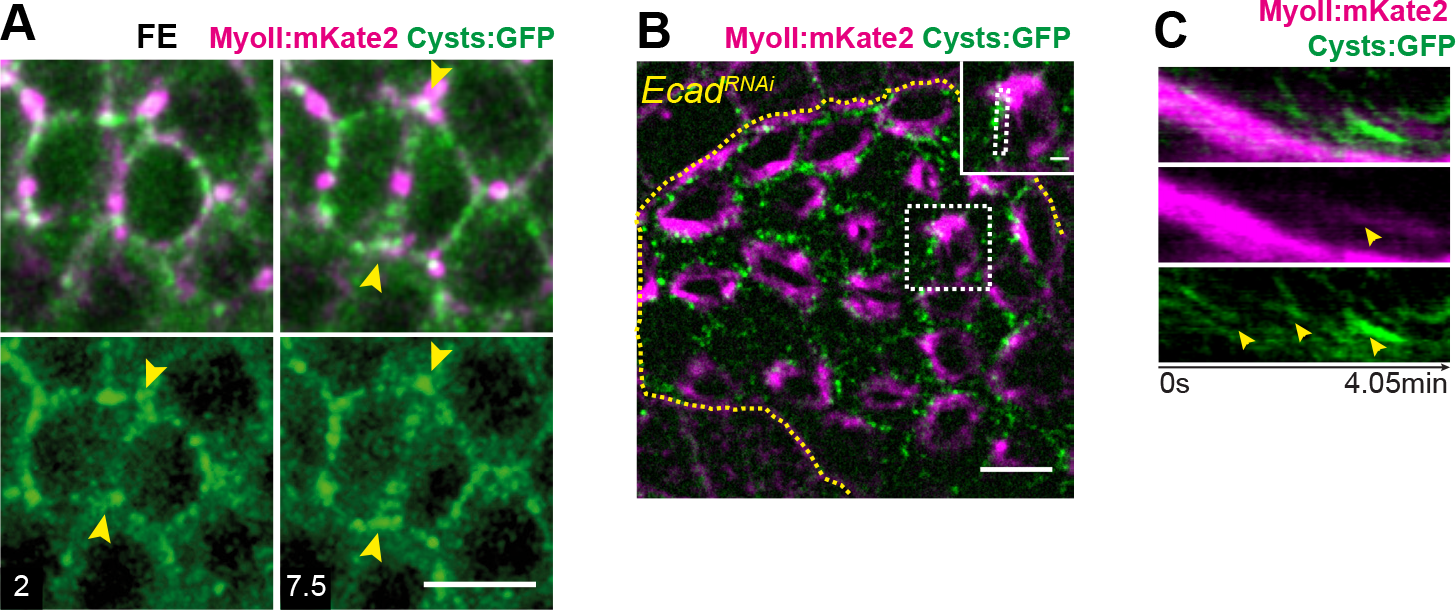
Analysis of Cysts localization and function, related to Figure 2. Unless otherwise indicated, all confocal images are apical top views at the level of the AJ in the notum, and time (min, indicated in the lower left image corner) is set to 0 at the onset of cytokinesis marked by the initial deformation of AJ by the constriction of the cytokinetic ring. **(A)** Time-lapse images of Cysts:GFP and MyoII:3xmKate2 during cytokinesis in the FE. Yellow arrowheads: Cysts:GFP accumulations at the rim of the cytokinetic ring. **(B)** Image of Cysts:GFP and MyoII:3xmKate2 in ctrl and *Ecad^RNAi^* interphasic cells. *Ecad^RNAi^* cells are marked by the expression of CAAX:tBFP (not shown), and a yellow dashed line outlines the boundary between ctrl and *Ecad^RNAi^* cells. Inset: close-up on the region outlined by the dashed white box. **(C)** Kymographs of Cysts:GFP (top and bottom) and MyoII:3xmKate2 (top and middle) within the region outlined in the inset in (B). Yellow arrowheads indicate the flows of MyoII:3xmKate2 and Cysts:GFP towards the medial apical region of the *Ecad^RNAi^* cell. Time (min) is set to 0 at the beginning of the kymograph. Scale bars: 5 µm (A, B), 1µm (inset in B).

**Figure S5:**
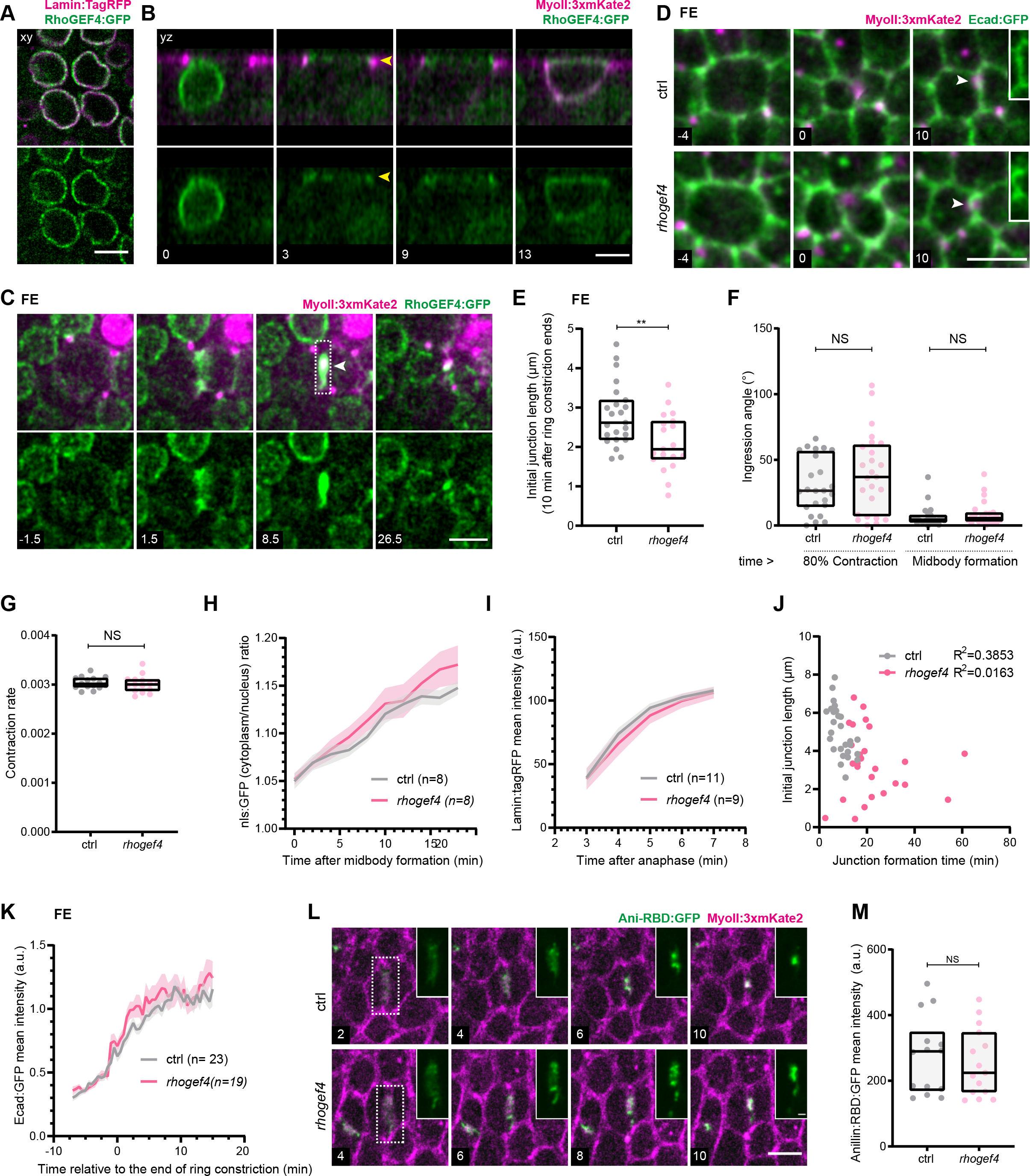
R**h**oGEF4 **localization and function in epithelial tissues, related to** Figure 3. Unless otherwise indicated, all confocal images are apical top views at the level of the AJ in the notum, and time (min, indicated in the lower left image corner) is set to 0 at the onset of cytokinesis marked by the initial deformation of AJ by the constriction of the cytokinetic ring. For details on quantifications, see STAR Methods. (A) xy images at the level of the nuclei of RhoGEF4:GFP (top and bottom) and Lamin:TagRFP (top) in interphase cells. (B) yz time-lapse images of RhoGEF4:GFP (top and bottom) and MyoII:3xmKate2 (top) from interphase to early cytokinesis. yz sections are shown at the level of the contractile ring. Time (min) is set to 0 at the start of the time-lapse sequence. Yellow arrowhead: AJ. (C) Time-lapse images of RhoGEF4:GFP (top and bottom) and MyoII:3xmKate2 (top) during cell division in the FE. White arrowhead: midbody. White dashed box: daughter cell interface. (D) Time-lapse images of Ecad:GFP and MyoII:3xmKate2 in ctrl and *rhogef4* dividing cells in the FE. Time is set to 0 at the end of ring contraction. White arrowhead: midbody. Inset: daughter cell interface for the Ecad:GFP channel. (E) Box plot of initial AJ length (median ± interquartile range) in ctrl and *rhogef4* dividing cells in the FE. Junction length measurements were performed 10 min after the end of ring constriction. (F) Box plot of daughter-daughter interface angle (median ± interquartile range) at 80% of apical ring contraction and at midbody formation in ctrl and *rhogef4* cells in the notum. (G) Box plot of ring contraction rate (median ± interquartile range) in ctrl and *rhogef4* cells in the notum. (H) Graph of the nls:GFP cytoplasmic/nuclear ratio (mean ± SEM) during late cytokinesis in ctrl and *rhogef4* cells in the notum. (I) Graph of Lamin:TagRFP levels (mean ± SEM) at the daughter cells nuclear envelope during cytokinesis in ctrl and *rhogef4* cells in the notum. (J) Graph of the initial AJ length as a function of the time of AJ formation in ctrl and *rhogef4* dividing cells in the notum. R^2^ values for linear regressions are indicated. (K) Graph of Ecad:GFP levels at the daughter cell interface (mean ± SEM) during *de novo* junction formation in ctrl and *rhogef4* cells in the FE. Time is set to 0 at the end of ring constriction. (L) Time-lapse images of MyoII:3xmKate2 and Ani-RBD:GFP for ctrl and *rhogef4* dividing cells during cytokinesis. Inset: close-up on the Ani-RBD:GFP signal at the apical contractile ring and midbody. (M) Box plot of Ani-RBD:GFP intensity (median ± interquartile range) at the apical contractile ring of ctrl or *rhogef4* cells in the notum. Scale bar: 5µm (*A, B, C, D, L)*, 1µm (inset in L). Mann-Whitney test: *NS*: non-significant. **: *p*<0.01

**Figure S6:**
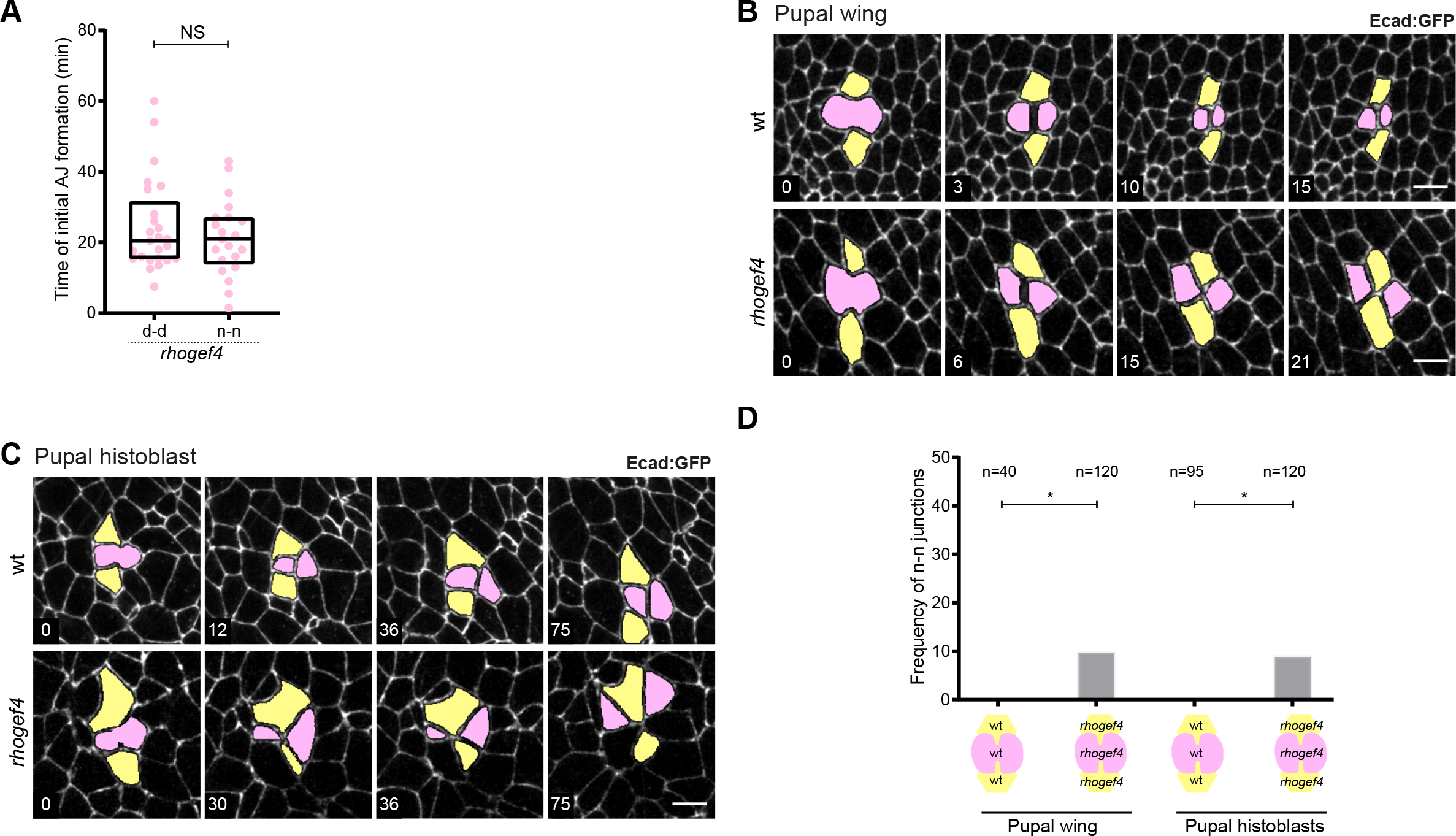
Cell topology upon cytokinesis in different epithelia, related to Figure 4. All images are apical top views at the level of the AJ labelled by Ecad:GFP. The dividing and daughter cells are coloured in pink and neighbouring cells in yellow. Unless otherwise indicated, time (min, indicated in the lower left image corner) is set to 0 at the onset of cytokinesis marked by the initial deformation of AJ by the constriction of the cytokinetic ring. (A) Graph of the timing of the initial AJ formation in *rhogef4* dividing cells at which a d-d or a n-n junction are initially formed, in the notum epithelium. The time (min) is set to 0 at midbody formation. (B) Time-lapse images of Ecad:GFP from cytokinesis onwards in wt or *rhogef4* animals in the pupal wing epithelium. (C) Time-lapse images of Ecad:GFP from cytokinesis onwards in wt or *rhogef4* animals in the pupal histoblast nest epithelium. (D) Graph of the frequency of total n-n junctions formed upon cell division in wt and *rhogef4* mutant animals in the pupal wing and histoblast epithelial tissues. Scale bar: 5µm. Mann-Whitney test, NS : non significant (A). Z score proportion test (D). *: p<0.05.

Movie S1: Dynamics of RhoGEF/GAP during epithelial cell division in the notum, related to Figure 1.

‘xy’ indicates apical confocal top view at the level of the AJ, while ‘yz’ denotes apical-basal confocal section at the level of the cytokinetic ring or the midbody, MyoII:3xmKate2 is shown in magenta. Time (min) is set to 0 at the onset of cytokinesis marked by the initial deformation of AJ by the constriction of the cytokinetic ring.

(**A**) yz time-lapse movie of Cdep:GFP and MyoII:3xmKate2 during cell division in the notum. (**B-I**) xy time-lapse movies of CdGAPr:GFP (B), Exn:GFP (C), Graf:GFP (D), Mbc:GFP (E), RhoGAP1A:GFP (F), RhoGAP5A:GFP (G), RtGEF:GFP (H) and Ziz:GFP (I) with

MyoII:3xmKate2 (B-I)) during cell division in the notum.

Scale bar: 5µm.

Movie S2: Dynamics of RhoGEF/GAP during epithelial cell division in the FE, related to Figure 1.

‘xy’ indicates apical confocal top view at the level of the AJ. Time (min) is set to 0 at the onset of cytokinesis marked by the initial deformation of AJ by the constriction of the cytokinetic ring.

(**A-F**) xy time-lapse movie of CG46491:GFP (A), Graf:GFP (B), Mbc:GFP (C), RhoGAP1A:GFP (D), RhoGAP5A:GFP (E) and RhoGAP15B:GFP (F) with MyoII:3xmKate2

(A-F) during cell division in the FE. Scale bar: 5µm.

Movie S3: Localisation and role of Cysts during epithelial cell division, related to Figure 2.

‘xy’ indicates confocal top view in the notum. Time (min) is set to 0 at the onset of cytokinesis marked by the initial deformation of AJ by the constriction of the cytokinetic ring.

(A) Time-lapse movie of Cysts:GFP and MyoII:3xmKate2 (top) in a dividing cell and its neighbours.

(B) Time-lapse movie of clonally expressed His2B:RFP in a dividing cell (left) and Cysts:GFP (left, right) in its neighbours. Cysts:GFP images are projections at the level of the cell apical surface while His2B:RFP are projections at the level of the chromosomes.

(C) Time-lapse movies of Ani-RBD:GFP and His2B:RFP in ctrl (left) and *cysts* (right) cells neighbouring a dividing cell. Ani-RBD:GFP and His2B:RFP are expressed clonally; *cysts* cells are marked by the loss of His2B:RFP signal in the right panels. Images are projections of 3.5µm from encompassing the apical surface and part of the nuclei.

Scale bar: 5µm.

Movie S4: Role of Cysts during epithelial cell division, related to Figure 2.

Time-lapse ‘xy’ movie of Ecad:GFP and MyoII:3xmKate2 in the context of a ctrl dividing cell surrounded by ctrl (bottom cell) and *cysts* (top cell) neighbouring cells. *cysts* cells are marked by the loss of nls:GFP signal.

Scale bar: 5µm.

**Movie S5: RhoGEF4 dynamics and function during epithelial division, related to** Figure 3. ‘xy’ indicates confocal top view in the notum. Time (min) is set to 0 at the onset of cytokinesis marked by the initial deformation of AJ by the constriction of the cytokinetic ring.

(A) Time-lapse movie of RhoGEF4:GFP (top and bottom) and MyoII:3xmKate2 (bottom) from interphase to late cytokinesis.

(B) Time-lapse movies of Ecad:GFP, MyoII:3xmKate2 and His2A:RFP in ctrl and *rhogef4* dividing cells. *rhogef4* cells are marked by the absence of His2A:RFP expression.

(C) Time-lapse movie of Ecad:3xmKate2 and Rac1:GFP in ctrl (left) and *rhogef4* (right) dividing cells during cell division.

(D) Time-lapse movie of Ecad:3xmKate2 and Utr-ABD:GFP in ctrl (left) and *rhogef4* (right) dividing cells during cell division.

Scale bar: 5µm.

Movie S6: *de novo* junction formation in control and *rhogef4* cells, related to Figure 4.

Time-lapse ‘xy’ movie of Ecad:GFP (left) and *rhogef4* (right) dividing cells during cytokinesis and junction formation. The dividing and daughter cells are coloured in pink and the neighbouring cells in yellow. Time (min) is set to 0 at the onset of cytokinesis marked by the initial deformation of AJ by the constriction of the cytokinetic ring.

Scale bar: 5µm.

Table S1: Plasmids, BAC and Oligonucleotides for tagged RhoGEF and RhoGAP as well as RMCE RhoGEF and RhoGAP alleles.

(**Column A-C**): RhoGEF and RhoGAP names, chromosomal position (B) and DNA strand (C).

(**Column D**): Closest human orthologues. We only report the best score Flybase hits for the human orthologues.

*For RhoGEF and GAP tagged at their endogenous locus by CRISPR/Cas9 mediated homologous recombination (first and second tabs, top)*.

(**Column E-G**): The RhoGEF/GAP were tagged with GFP or mKate2 at their endogenous locus in C-terminal or N-terminal using CRISPR/Cas9 mediated homologous recombination. GFP:xxx or mKate2:xxx versus xxx:GFP or xxx:mKate2 indicate that the GFP or mKate2 tag were inserted in N-terminal versus C-terminal. The tagged alleles generated are listed in (E) and the other available HR donor plasmids in (F). The precise positions of the GFP or mKate2 tags are given in single letter aa code, and the tagged isoforms are indicated (G). Tag insertion can be associated with a deletion of a few aa as well as deletion of non-coding sequences.

(**Column H-O**): The parental donor vectors (H) used for the cloning of the homologous region 1 (I-L) and homologous region 2 (M-P). The genomic start (I, M) and end (J, N) positions of each homologous region, as well as the forward (K, O) and reversed primer (L, N) used for PCR and subsequent cloning in the parental donor vector. Nucleotides indicated in red are the regions matching the parental donor plasmid sequence used for cloning by SLIC.

(**Column P-Y**): The parental guide vector (P) used for the cloning of the CRISPR/Cas9 RNA expressing plasmids. We used 2 distinct gRNA either cloned in two different pCFD3- dU6:3gRNA parental plasmid or cloned in the same multiplex gRNA pCFD5 (Addgene

#73914) plasmid. The genomic start (R,V) and end (S,W) positions of the two guides as well as the forward (T,X) and reverse (U,Y) primers used for cloning. When using the pCFD3- dU6:3gRNA (Addgene #49410), the forward and reverse oligonucleotides were annealed and cloned in the pCFD3-dU6:3gRNA. When using the pCFD5 plasmid (Addgene #73914), the forward and reverse oligonucleotides were used to PCR amplify the sequences containing the 2 guides, and the GlytRNA sequence from the parental plasmids. Nucleotides indicated in red are the regions matching the plasmid sequence used for cloning.

(**Column Z**): Flybase link.

*For RhoGEF and GAP tagged using BAC recombineering (first and second tabs, bottom)*.

(**Column E-G**): The RhoGEF and GAP BAC were tagged with GFP at their C-terminal or N-terminal. GFP:xxx versus xxx:GFP indicate that the GFP tag is inserted in N-terminal versus C-terminal. The available transgenic BAC lines and BAC with a GFP insertion are indicated in E and F respectively. The position of the GFP tag is given in single letter aa code, and the tagged isoforms are indicated (G).

(**Column H-O**): The parental BAC and its size (H) used for inserting the GFP cassette. The BAC nucleotide positions of the homologous regions (I,L) and the forward and reversed primers (M,N) sequences used to PCR amplify the HR regions with the GFP-lox-neo-lox cassette. Nucleotides indicated in red are the regions homologous to GFP-lox-neo-lox cassette.

(**Column P**): Docking site used for transgenesis of the tagged RhoGEF or RhoGAP BAC. (**Column Q**): Flybase link.

*For RhoGEF and GAP RMCE allele (third and fourth tabs)*.

(**Column D,E**): Genomic nucleotide and aa positions of the expected deletions generated using CRISPR/Cas9 mediated homologous recombination. The aa are given on the coding sequence of the longest isoform, the length of which is indicated. Isoforms that will be affected (F).

(**Column F-N**): The parental donor vectors (F) used for the cloning of the homologous region 1 (G-J) and homologous region 2 (K-N). The genomic start (G, K) and end (H, L) positions of each homologous region as well as the forward (I, M) and reversed primers (J, R) used to PCR amplify the HR. Nucleotides indicated in red are the ones matching the parental plasmid sequence used for SLIC.

(**Column O-W**): The parental guide vector (O) used for the cloning of the N-terminal CRISPR/Cas9 RNA expressing guides. The genomic start (Q,U) and end (R,V) positions of the two guides as well as the forward (S,W) and reverse (T,X) primers used for cloning. When using the pCFD5 (Addgene #73914) plasmid, the two guides were inserted in the forward and reverse primers. Nucleotides indicated in red are the regions matching the plasmid sequence used for cloning.

(**Column P-AF**): The parental guide vector (P) used for the cloning of the CRISPR/Cas9 RNA expressing plasmids. We used 4 distinct gRNA (2 in Nter and 2 in Cter) cloned in 4 different pCFD3-dU6:3gRNA parental plasmids, 4 distinct gRNA (2 in Nter and 2 in Cter) cloned in 2 multiplex gRNA pCFD5 (Addgene #73914) plasmids or 2 distinct gRNA (1 in Nter and 1 in Cter) cloned in the same multiplex gRNA pCFD5 (Addgene #73914) plasmid. The genomic start (P,T,Y,AC) and end (Q,U,Z,AD) positions of the guides, as well as the forward (R,V,AA,AE) and reverse (S,W,AB,AF) primers used for cloning. When using the pCFD3- dU6:3gRNA (Addgene #49410), the forward and reverse oligonucleotides were annealed and cloned in the pCFD3-dU6:3gRNA. When using the pCFD5 plasmid (Addgene #73914), the forward and reverse oligonucleotides were used to PCR amplify the sequences containing the 2 guides and the GlytRNA sequence from the parental plasmids. Nucleotides indicated in red are the regions matching the plasmid sequence used for cloning.

Table S2: RhoGEF/GAP localizations in interphase and division in the notum and the FE.

*Tagged RhoGEF (Tab 1) and RhoGAP (Tab 2)*.

(**Column A**): RhoGEF or RhoGAP tagged protein (A), GFP:xxx or mKate2:xxx versus xxx:GFP or xxx:mKate2 indicate that the GFP or mKate2 tag is inserted in N-terminal versus C-terminal (A).

(**Column B-D):** Signal intensity (B), localization in interphase (C) and during cell division (D) in the notum. Signal intensity is given in shades of green. Not detected: RhoGEF/GAP with no detectable signal. NA: not applicable. *: RhoGEF/GAP with low signal to noise for which only still images were acquired during cell division.

(**Column E-G):** Signal intensity (E), localization in interphase (F) and during cell division (G) in the FE. Signal intensity is given in shades of green. Not detected: RhoGEF/GAP with no detectable signal. NA: not applicable.

(**Column H):** Closest human orthologues. The best scored Flybase hits for the human homologues are listed.

(**Column I**) Link to Flybase webpage for each gene.

## STAR METHODS

`o` KEY RESOURCES TABLE
`o` RESOURCE AVAILABILITY

`o` Lead contact
`o` Materials availability
`o` Data and code availability
`o` EXPERIMENTAL MODEL AND SUBJECT DETAILS

`o` Fly husbandry and stocks
`o` METHOD DETAILS

`o` Molecular biology and transgenesis
`o` Transgenic Fluorescently tagged RhoGEF/GAP Library
`o` Plasmid Library for generating RhoGEF/GAP deletion RMCE alleles
`o` Genetics, somatic mutant clones and RNA interference
`o` Pupa and egg chamber mounting for live imaging
`o` Microscopy
`o` Localization screen of tagged RhoGEF/GAP in live epithelial tissues
`o` Optogenetics
`o` QUANTIFICATION AND STATISTICAL ANALYSIS

`o` Dynamics of cytokinesis, membrane juxtaposition and AJ formation
`o` Fluorescent reporter levels
`o` Figure preparation
`o` Sampling and Statistics

## RESOURCE AVAILABILITY

### Lead contact

Further information and requests for resources and reagents should be directed to, and will be fulfilled by the lead contact, Yohanns Bellaïche.

### Materials availability

Flies and plasmids are available from the lead contact.

### Data availability

No large-scale datasets have been generated in this study. The data that support the findings of this study are available from the corresponding authors.

### Code availability

Fiji customized Macros available upon request.

## EXPERIMENTAL MODEL AND SUBJECT DETAILS

### Fly husbandry and stocks

Flies were grown on standard molasses/cornmeal/yeast food at 18°C or 25°C, and experiments were performed at 25° unless otherwise specified. *Drosophila* melanogaster stocks used in this study and associated references are listed in the Key Resource Table.

## METHOD DETAILS

### Molecular Biology and Transgenesis

Table S1 lists BAC and plasmids generated in this study, their parental BAC or plasmids, as well as the oligonucleotides used. Oligonucleotides were bought from Sigma. Cloning was performed by primer annealing and ligation, or by SLIC (Li and Elledge, 2012). All regions amplified by PCR as well as junctions between tag and genomic sequences were checked by sequencing. Transgenesis was performed by BestGene or Rainbow transgenesis services.

### Transgenic Fluorescently tagged RhoGEF/GAP Library

#### Recombineering based Tagging

To create the GFP-tagged transgenes of CG15611, CG43658, RhoGEF2, RtGEF, Trio, RacGAP84C, RhoGAP100F, RhoGAP102A and RhoGAP93B expressed under the control of their endogenous promoter, we used recombineering (Venken et al., 2009) to introduce an in-frame GFP sequence in N- or C-term of the open reading frame in BAC genomic clones (see Table S1 for GFP position, tagged isoforms and BAC clones from the BACPAC Resources Center (Venken et al., 2009)). To insert a GFP sequence at the amino acid positions indicated in Table S1 by recombineering, a GFP sequence and a neomycin resistance cassette flanked by loxP sites and homologous sequences were amplified by PCR and recombined in each BAC (Venken et al., 2008). Upon neomycin selection, the cassette was removed by Cre-mediated recombination leaving behind a 78 bp loxP site sequence. Each GFP insertion was verified by sequencing. Each BAC construct was integrated at the PBac{y[+]- attP-9A}VK00033 landing site at 65B2 and their genomic insertion was confirmed by PCR and sequencing.

#### CRISPR/Cas9 mediated tagging

The remaining RhoGEF/GAP were tagged with either GFP or mKate2 at their endogenous locus using CRISPR/Cas9 mediated homologous recombination (Kanca et al., 2019; Port et al., 2014). To generate the tagged alleles by CRISPR/Cas9-mediated homologous recombination, guide RNA sequences were selected using CRISPR Optimal Target Finder (http://targetfinder.flycrispr.neuro.brown.edu/index.php, (Gratz et al., 2014)), and corresponding oligonucleotides were cloned into the pCFD5 (Addgene #73914) or pCFD3- dU6:3gRNA (Addgene #49410) guide vectors (Table S1). Homology sequences were cloned into homologous recombination vectors harbouring a *hs-miniwhite* cassette flanked by two loxP sites (Huang et al., 2008; Pinheiro et al., 2017) and an N- or C-terminal GFP or mKate2 sequences using the pCRISPR-GFP-Nter, pCRISPR-GFP-Cter, pCRISPR-mKate-Nter and pCRISPR-mKate-Cter vectors (Pinheiro et al., 2017). The two homologous regions were cloned using the primers indicated in Table S1. Plasmids expressing the guide RNAs for each RhoGEF/GAP tagging were co-injected in *vas-cas9* embryos. When injecting donors for RhoGEF/GAP for which both the mKate2 and GFP donor plasmids were cloned, we co-injected GFP and mKate2 donor plasmids at a 2/3 and 1/3 ratio to favour GFP tagging. Upon excision of the *mini-white* (*mw*) transgenesis marker using hs-Cre lines on chromosome II or III, each fluorescently tagged allele was verified by sequencing.

### Plasmid Library for generating RhoGEF/GAP deletion RMCE alleles

To facilitate the generation of loss of function alleles of each RhoGEF/GAP using CRISPR/Cas9 mediated homologous recombination, we generated a plasmid library of guide and donor plasmids to replace most of each RhoGEF/GAP coding region by a *mw* gene flanked by 2 attP Phi31 recombination sites (See Table S1 for guide and donor plasmids and deletions positions) using the pCRISPR-del (Pinheiro et al., 2017) or pCRISPR-del-HighThrouput (Kanca et al., 2019) plasmids. Using the well-established approach known as RCME based on the ϕ31 recombinase (Fig. S1B, (Zhang et al., 2014)), one can then efficiently replace the *mini-white* gene by any DNA sequence to perform structure-function analysis or tagging with any fluorescent or non-fluorescent tags (Fig. S1B). The plasmid library was used to generate *cyst^ΔRMCE^* and *rhogef4^ΔRMCE^* null alleles. *cyst^ΔRMCE^* is associated with a deletion of the *cysts* reading frame from G233 to F1309 (numbered according to the isoform RA, 1309 aa) and *rhogef4^ΔRMCE^* deletes from M1 to F647 (numbered according to the isoform RA, 647 aa) of the *rhogef4* reading frame.

### Genetics, somatic mutant clones and RNA interference

Both female and male animals were used in experiments in the pupa, except in shown in Fig. 2E for which females were used; adult females were used for experiments in the FE. Experiments using the Gal4/UAS, the Gal4/Gal80^ts^/UAS systems (Brand and Perrimon, 1993; McGuire et al., 2003), hs-FLP induced FLPout clones and FRT site mitotic recombination (Golic and Lindquist, 1989) were done as previously described (Bosveld et al., 2012). *cyst^ΔRMCE^*,

*rhogef4^ΔRMCE^*, *Rho1^72M1^* and GFP:RhoGEF4 somatic mutant clones were generated by FRT/FLP induced mitotic recombination, and mutant cells were marked by the loss of fluorescent markers. Clones in the pupa were generated via a 1h heat-shock at 37°C 3-4 days before live-imaging. Clones in the FE were induced via heat-shocking 2 times for 2h at 37°C 2- 5 days before imaging. The *cysts,rhogef4* double mutant cells were obtained by inducing *cyst^ΔRMCE^* clones in *rhogef4^ΔRMCE^* mutant animals. Experiments in the pupal wing (Fig. S6B,D) were performed using the *rhogef4^ΔRMCE^*/Df(3L)BSC117 genotype; Df(3L) deletes the *rhogef4* locus. Experiments in Fig. S5D, S5E and S5K were done in ovaries dissected from *rhogef4^ΔRMCE^* homozygous flies expressing Ecad:GFP and MyoII:3xmKate2, and Ecad:GFP, MyoII:3xmKate2 flies were used as control.

The *pnut^RNAi^, ecad^RNAi^ or rok^RNAi^* clones were generated by driving the expression of *UAS- pnut^dsRNA^*, *UAS-ecad^dsRNA^* or *UAS-rok^dsRNA^*, respectively, using hs-FLP induced flp-out on the Act>Cdc2>Gal4, UAS-CAAX:tBFP line to express Gal4 and label the clones by membrane localized tBFP.

### Pupa and egg chamber mounting for imaging

Pupae were collected and mounted for dorsal thorax, wing or histoblast live imaging as described previously (Bardet et al., 2013; Bosveld et al., 2012; Pinheiro et al., 2017). Dissected egg chambers were imaged live in culture medium (Schneider’s medium, Sigma-Aldrich) supplemented with 10% Fetal bovine serum (Thermo Fisher) and 200 μg/uL insulin (Sigma-Aldrich) as previously described (Osswald et al., 2022). For imaging of fixed tissue, egg chambers were fixed in 4% paraformaldehyde solution in PBT (PBS + 0.2% Tween 20, Sigma-Aldrich) for 20 min, washed with PBT and mounted with Vectashield with DAPI (Vector Laboratories).

### Microscopy

Microscopes, software and imaging: Samples were imaged at 25°C, unless otherwise specified.

#### Imaging of pupal tissues

The imaging was performed on: (i) an inverted confocal spinning disk microscope CSU-W1 (Andor/Roper/Nikon) equipped with a sCMOS camera (Flash4 Hamamatsu), and a Borealis module from Andor for better field homogeneity, using either a 60x (NA1.4 OIL PL APO L) or 100x (NA1.45 OIL PL APO L); (ii) an inverted confocal spinning disk microscope from Zeiss (CSUW1, Roper/Zeiss) equipped with a sCMOS camera (Flash4 Hamamatsu), a Visitron module for better field homogeneity, using either 60x or 100x NA1.4 OIL DIC N2 PL APO VC objectives; (iii) Inverted spinning disk wide confocal microscope CSU-W1 (Roper/Nikon/GATACA) equipped with a sCMOS camera (BSI camera), a field illumination homogenizer, using either 60x or 100X (NA 1.4 OIL N2 PL APO VC) objectives. The Metamorph autofocus function was used for some time-lapse acquisitions.

#### Imaging of egg chambers

The imaging was done on either: (i) an Andor XD Revolution Spinning Disk Confocal system (Yokogawa CSU-22 unit) equipped with two solid state lasers (488nm and 561nm), an iXonEM+ DU-897 EMCCD camera, and built on an inverted Olympus IX81 microscope with a 60x (NA1.42 OIL PLAPON) or a 100x (NA1.40 OIL UPLSAPO) objective using iQ software (Andor); (ii) an inverted laser scanning confocal microscope Leica

SP8 (Leica Microsystems) with a 63x (NA1.30 GLYCEROL) objective; (iii) an inverted laser scanning confocal microscope Leica TCS SP5 II (Leica Microsystems) with a 63x (NA1.3 GLYCEROL) objective.

### Localization screen of tagged RhoGEF/GAP in live epithelial tissues

To determine the localization of GFP-tagged RhoGEF/GAP in living epithelial notum tissue, a series of sequential acquisitions were performed for each tagged protein. First, using MyoII:3xmKate2 as a reference, the GFP signal was acquired at different apical-basal levels both in interphase and dividing cells. Following this initial acquisition, two complementary acquisitions were used to better capture each RhoGEF/GAP localization or resolve their dynamics at a relevant position along the apical-basal axis: i) a single z-stack acquisition at high laser power for RhoGEF/GAP expressed at intermediate/low levels; ii) a z-stack and time-lapse serie using a time interval ranging from 30s to 4min, adjusted to reduce bleaching of the GFP signal, and to image cell division. In both cases, the apical and basal positions of the z-stack acquisitions were set to the relevant z-plans as determined by the initial acquisition.

The localizations of fluorescently tagged RhoGEF/GAP were determined in live and fixed egg chambers through the acquisitions of z-stacks (z-step between 0.5 and 1 μm): (i) at the surface of the egg chamber to cross-section the FE along the apical-basal axis (labelled as “xy” in figures); (ii) along the midsagittal region of the egg chamber, which enables the visualization of the whole apical-basal axis of the epithelium in a confocal section plane (labelled as “yz” in figures). MyoII:3xmKate2 or CellMask Orange Plasma membrane Stain (ThermoFisher; C10045; diluted 1:10 000 in *ex vivo* culture medium (Schneider’s medium (Sigma-Aldrich) supplemented with 10% Fetal bovine serum (Thermo Fisher) and 200 μg/uL insulin (Sigma-Aldrich) and washed twice with *ex vivo* culture medium before imaging) were used to label the actomyosin cytoskeleton or the plasma membrane, respectively, to resolve the fluorescently tagged RhoGEF/GAP position along the apical-basal axis.

### Optogenetics

Experimental crosses were protected from light by wrapping the Drosophila tubes into an aluminium foil and were grown at 18°C. White pupae (0 hAPF) were selected under white light, and then kept in the dark until the experiment performed at 29°C to induce the expression of the LARIAT transgene. Pupae were mounted under amber light (white light lamp source equipped with a 630/692nm BrightLine® single-band bandpass filter) in a dark room to avoid LARIAT activation. Photoactivation of LARIAT was achieved using repeated illumination (6 0.5µm z-section every 30 sec) with a 488nm laser set at 20-50% of its maximal power on a spinning disk microscope. This routine was selected as it promotes efficient Ecad:GFP clustering at the AJ.

## QUANTIFICATIONS AND STATISTICAL ANALYSIS

Quantifications were performed using the Fiji software and a set of home-made macros.

### Dynamics of cytokinesis, membrane juxtaposition and AJ formation

#### Constriction rate and time of 80% contraction

The rates of constriction were determined as in (Herszterg et al., 2013). Briefly, the speed of cytokinetic ring constriction in the plane of the AJ was used as a proxy of the cytokinetic constriction rate on MyoII:3xmKate2 and Ecad:GFP confocal time lapse images. To determine constriction rate and the time-point corresponding to 80% of constriction, manual measurements of the initial apical cell length at the site of ring formation, and of the length of apical contractile ring were performed using a home-made Fiji macro. The rate of constriction was determined as the slope of the linear fit of the contractile ring length, normalized to its initial apical length at the onset of cytokinesis, as a function of time (Herszterg et al., 2013).

#### Angle formed by the ingressing AJ

In control and mutant conditions, the angle (from 0° to 90°) formed by the ingressing AJ was quantified in time-lapse movies of Ecad:GFP and MyoII:3xmKate2, as schematically represented in Figure 2H; a 0° angle corresponds to maximal daughter cell membrane juxtaposition. The angles were measured manually at 80% of constriction of the dividing cell using a Fiji Macro as in (Pinheiro et al., 2017).

#### Junction formation dynamics in the notum

The timing and the initial length of the *de novo* AJ formed upon division were determined in Ecad:GFP and MyoII:3xmKate2 time-lapse movies. The time of *de novo* AJ formation was calculated relative to the timing of midbody formation as the timepoint at which Ecad:GFP signal is observed along the whole d-d or n-n interface. The initial AJ length was measured for d-d interface at the time of *de novo* AJ formation.

#### Junction formation dynamics in the FE

Both Ecad:GFP signal at the *de novo* AJ formed upon cell division and junctional length were determined from time-lapse movies of z-stacks acquired at the surface of Ecad:GFP and MyoII:3xmKate2 egg chambers from control and *rhogef4^ΔRMCE^* mutant animals. As the egg chamber curvature prevents the visualization of the whole FE junctional network in individual planes, an adapted projection function (Epitools in Matlab (Heller et al., 2016)) was used to project the Ecad:GFP and MyoII:3xmKate2 signals from the whole junctional plane into a 2D image. The Ecad:GFP signal at the new cell interface and the interfaces of interphasic cells (used as a reference for normalization) as well as the background signal were manually tracked throughout the time-lapse movie to determine the dynamics of Ecad:GFP fluorescence intensity at the new interface during cell division. To quantify the initial junction length, the length of the newly-formed interface was measured 10 minutes after ring constriction. To take egg chamber curvature into consideration in this analysis, initial junction length was measured using the original z-stacks.

#### Neighbouring membrane withdrawal

The analysis of neighbouring cell membrane withdrawal duration was determined at the interface between PH:GFP and PH:mCherry cell patches to distinguish the membranes of the dividing and daughter cell membranes from those of the neighbouring cells, focusing on the apical cell level (Herszterg et al., 2013). The duration of neighbouring membrane withdrawal was calculated as the difference between the timepoint of full neighbouring membrane insertion and full membrane withdrawal from the dividing cells.

#### Frequency of rearrangements upon cytokinesis

The analyses of the frequency of n-n AJ formation upon cytokinesis were performed on time-lapse movies of Ecad:GFP in wt and *rhogef4^ΔRMCE^* or *rhogef4^ΔRMCE^* /Df (3L)BSC117 animals, as well as in *cysts ^ΔRMCE^* clones neighbouring dividing cells in wt or *rhogef4^ΔRMCE^* tissues. The criterion for the formation of a n-n AJ was the formation of a n-n Ecad positive interface for at least one time-point. The formation of n-n AJ occurs either at the time of the initial AJ formation upon cytokinesis, or immediately after the formation of a very short d-d or a 4-fold cell vertex. The formation of n-n AJ formation upon cytokinesis was also observed in *rhogef4^ΔRMCE^* clones.

### Fluorescent reporter signal levels

#### MyoII, Ani-RBD and Cysts accumulations in neighbouring cells

In control and mutant conditions, the MyoII accumulations in the neighbours were quantified in time-lapse movies of Ecad:GFP and MyoII:3xmKate2 at 80% of constriction of the dividing cell, as previously performed (Pinheiro et al., 2017). Upon determination of the time-point corresponding to 80% of the constriction (see above), the accumulation of MyoII:3xmKate2 was quantified as the average of the 2 time points closest to the 80% ring constriction time point. To quantify MyoII:3xmKate2 accumulation in an unbiased manner using Fiji, the mean MyoII:3xmKate2 intensity of the neighbouring cells in each frame was used to threshold the image and select the pixels above the mean intensity; thus, defining a ROI corresponding to the MyoII:3xmKate2 accumulation at the base of the ingressing AJ. MyoII accumulation in the neighbouring cells was determined as the integrated density of MyoII:3xmKate2 in the ROI normalized by the mean MyoII:3xmKate2 cortical intensity in the neighbours.

The accumulation of Ani-RBD:GFP in cells neighbouring a control dividing cell was analysed in time-lapses movies of His2B-RFP and Ani-RBD:GFP, and visually compared between control and cysts neighbours. Ani-RBD:GFP and His2B:RFP are expressed clonally; *cysts* cells are marked by the loss of His2B:RFP signal in the bottom panels. Ani-RBD:GFP was analysed in cells adjacent to a cell that divides roughly parallel to them and that is devoid of Ani- RBD:GFP signal.

The localization of Cysts:GFP during cytokinesis was analysed in time-lapse movies of Cysts:GFP together with Ecad:3xmKate2 or MyoII:3xmKate2.To assess the effect of loss of *pnut* knock-down on Cysts accumulation in neighbouring cells, the Cysts:GFP signal at the rim of the cytokinesis ring was compared in cells neighbouring wt or *pnut^RNAi^* (marked by the expression of CAAX:tBFP) dividing cells in Cysts:GFP, Ecad:3xmKate2 time-lapse movies. Using ctrl cells, we first defined that Cysts:GFP accumulation usually begins 3min after the initial deformation of the apical cell junctions due to ring contraction and it lasts for 4min. Within this time window, we then visually compared Cysts:GFP signal at the rim of cytokinesis ring in cells neighbouring wt and *pnut^RNAi^* dividing cells.

To assess the effect of *rok* knock-down on Cysts accumulation in the neighbouring cells during cytokinesis, the accumulation of Cysts:GFP at the rim of cytokinesis was evaluated in ctrl and *rok^RNAi^* (marked by the expression of CAAX:tBFP) neighbours cells surrounding a ctrl dividing cell in Cysts:GFP, MyoII:3xmKate2 time-lapse movies. The accumulation of Cysts:GFP was compared in the *rok^RNAi^* neighbours (*rok^RNAi^* side) and ctrl cells (ctrl side) surrounding the same dividing cells.

#### Accumulation of Fluorescent reporters in the contractile ring and daughter cell interface

The analyses of Ani-RBD:GFP, Pak3-RBD:GFP, Scar:GFP, Rac1:GFP and Utr-ABD:GFP signal in the dividing cell were performed on time-lapse movies, together with Ecad:3xmKate2 or MyoII:3xmKate. The mean intensity of each GFP signal was measured at the apical surface level by manually drawing a region of interest (ROI). The measurements were then averaged over several time-points. The average measurements were corrected by subtracting the averaged cytoplasmic signal in anaphase or cytokinesis. The ROI and timing of measurements were set based on the localization of the GFP and its temporal dynamics: Ani-RBD:GFP was measured at the apical contractile ring starting 2 min after the beginning of cytokinesis (marked by the initial deformation of the apical AJ) and averaged over 6 min (i.e., 3 time-points) ; Pak3- RBD:GFP and Scar:GFP were quantified at the daughter cell interface starting at the time of midbody formation and averaged over 10 and 6 min (i.e., 5 and 3 time-points), respectively. Rac1:GFP and Utr-ABD:GFP were measured at the daughter cell interface starting at the time of Ecad detachment from contractile ring and averaged over 8 and 10 min (i.e., 4 and 10 time- points), respectively.

#### Lamin:TagRFP and nls:GFP in telophase daughter cell nuclei

The analysis of Lamin:TagRFP intensity in daughter cell nuclei was performed in time-lapse movies of Lamin:TagRFP and Ecad:GFP. The Lamin:TagRFP intensity was measured at the nuclear envelope upon nuclear reformation during cytokinesis, starting 3 min after the beginning of anaphase during 5 min (i.e., 5 timepoints). To evaluate if nuclear envelope reformation was affected by RhoGEF4 loss of function, we determined nls:GFP reincorporation to the nuclei since defective nuclear sealing reduces nuclear localization of nls:GFP reporter (Olmos et al., 2016). The analyses were performed in nls:GFP, EcadGFP and MyoII:3xmKate2 time-lapse movies in wt and *rhogef4^ΔRMCE^* conditions. The nls:GFP mean intensity was measured in the cytoplasm and inside the reforming nuclei for 20 min starting from midbody formation. At this time, a weak nls:GFP signal allows to start distinguishing the nuclei. The nuclear/cytoplasmic ratios were then calculated for each cell over time.

#### Ecad and Cysts colocalization upon LARIAT photoactivation

To evaluate the hypothesis that a decrease in Ecad levels mediated by LARIAT will induce Cysts recruitment, we used notum tissues expressing Ecad:GFP, Cysts:mKate and LARIAT, and photoactivated by blue light illumination. On those tissues, we measured Cysts:mKate and Ecad:GFP levels in junctional regions encompassing Ecad gaps, or in junctional regions without gaps as internal ctrl. For this, we randomly selected each type of region, manually drew a line (1.3-2.6µm length) on each region of interest, and measured the profile of each signal using the

Plot profile function in Fiji. For each single line profile of Cysts:mKate-Ecad:GFP, we calculated Spearman correlation coefficients using Graphpad.

### Figure preparation

Images were subjected to bleach correction, brightness and contrast modifications and rotation using Fiji. Image denoising was performed using the Fiji despeckle filter, the Gaussian blur Fiji filter, or using a home-made denoising macro in Fiji. When necessary, time-lapse stacks were corrected for apical-basal z-drifts using a custom Fiji macro, or for xy-drifts using the MultistackReg Fiji plugin. For xy views in the notum, all images shown are maximal or sum projections of the relevant z-sections, whereas in yz views all images shown are sum projections of the relevant apical-basal transverse section generated using the Fiji stack reslice tool. As the egg chamber curvature prevents the visualization of the whole FE junctional network in individual planes, an adapted projection function (Epitools in Matlab (Heller et al., 2016)) was used to project the Ecad:GFP and MyoII:3xmKate2 signals from the whole junctional plane into a 2D image. For FE data, both xy and yz views are maximal projections of the relevant z-sections taken, respectively, at the surface or the midsagittal region of the egg chamber. Kymographs were generated using the multikymograph tool in Fiji.

Graphs were made using GraphPad (Prism). In the box plots, the box extends from the 25^th^ to 75^th^ percentiles (interquartile range), and the line in the middle of the box correspond to the median value. In graphs of any given quantity as a function of time, error bars correspond to the standard error of the mean (SEM).

### Sampling and Statistics

Sample sizes vary in each experiment. The experiments were repeated at least two independent times on a minimum of 3 animals. To determine the statistical difference between sets of continuous data, we performed non-parametric Mann-Whitney tests on GraphPad. To determine the statistical difference between two proportions, we performed two-sample Z-score tests for proportions (z score corresponding to 95% confidence interval = 1.96). For the LARIAT experiment, Spearman coefficients on line measurements of Ecad:GFP and Cysts:mKate2 intensities were calculated using Graphpad.

Significance is indicated by asterisks: *, *p*<0.05; **, *p*<0.01; ***, *p*<0.001; **** and *p*≤0.0001.

## REFERENCES

1. Abreu-Blanco, M.T., Verboon, J.M., and Parkhurst, S.M. (2014). Coordination of Rho Family GTPase Activities to Orchestrate Cytoskeleton Responses during Cell Wound Repair. Curr. Biol. 24, 144–155.

2. Acharya, B.R., Nestor-Bergmann, A., Liang, X., Gupta, S., Duszyc, K., Gauquelin, E., Gomez, G.A., Budnar, S., Marcq, P., Jensen, O.E., et al. (2018). A Mechanosensitive RhoA Pathway that Protects Epithelia against Acute Tensile Stress. Dev. Cell 47, 439–452.e6.

3. Aguilar-Aragon, M., Bonello, T.T., Bell, G.P., Fletcher, G.C., and Thompson, B.J. (2020). Adherens junction remodelling during mitotic rounding of pseudostratified epithelial cells. EMBO Rep. 21, e49700.

4. Aigouy, B., Farhadifar, R., Staple, D.B., Sagner, A., Röper, J.C., Jülicher, F., and Eaton, S. (2010). Cell Flow Reorients the Axis of Planar Polarity in the Wing Epithelium of Drosophila. Cell 142, 773–786.

5. Airoldi, S.J., McLean, P.F., Shimada, Y., and Cooley, L. (2011). Intercellular protein movement in syncytial Drosophila follicle cells. J. Cell Sci. 124, 4077–4086.

6. Arnold, T.R., Stephenson, R.E., and Miller, A.L. (2017). Rho GTPases and actomyosin: Partners in regulating epithelial cell-cell junction structure and function. Exp. Cell Res. 358, 20–30.

7. Bagci, H., Sriskandarajah, N., Robert, A., Boulais, J., Elkholi, I.E., Tran, V., Lin, Z.Y., Thibault, M.P., Dubé, N., Faubert, D., et al. (2020). Mapping the proximity interaction network of the Rho-family GTPases reveals signalling pathways and regulatory mechanisms. Nat. Cell Biol. 22, 120–134.

8. Bardet, P.L., Guirao, B., Paoletti, C., Serman, F., Léopold, V., Bosveld, F., Goya, Yû., Mirouse, V., Graner, F., and Bellaïche, Y. (2013). PTEN Controls Junction Lengthening and Stability during Cell Rearrangement in Epithelial Tissue. Dev. Cell 25, 534–546.

9. Bischoff, M., and Cseresnyes, Z. (2009). Cell rearrangements, cell divisions and cell death in a migrating epithelial sheet in the abdomen of Drosophila. Development 136, 2403–2411.

10. Bosveld, F., Bonnet, I., Guirao, B., Tlili, S., Wang, Z., Petitalot, A., Marchand, R., Bardet, P.- L., Marcq, P., Graner, F., et al. (2012). Mechanical Control of Morphogenesis by Fat/Dachsous/Four-Jointed Planar Cell Polarity Pathway. Science (80-.). 336, 724–727.

11. Brand, A.H., and Perrimon, N. (1993). Targeted gene expression as a means of altering cell fates and generating dominant phenotypes. Development 118, 401–415.

12. Buckley, C.E., and St Johnston, D. (2022). Apical–basal polarity and the control of epithelial form and function. Nat. Rev. Mol. Cell Biol. 23, 559–577.

13. Cadart, C., Zlotek-Zlotkiewicz, E., Le Berre, M., Piel, M., and Matthews, H.K. (2014). Exploring the function of cell shape and size during mitosis. Dev. Cell 29, 159–169.

14. Chaigne, A., and Brunet, T. (2022). Incomplete abscission and cytoplasmic bridges in the evolution of eukaryotic multicellularity. Curr. Biol. 32, R385–R397.

15. Chen, D.Y., Crest, J., Streichan, S.J., and Bilder, D. (2019). Extracellular matrix stiffness cues junctional remodeling for 3D tissue elongation. Nat. Commun. 10, 1–15.

16. Chu, Y.S., Thomas, W.A., Eder, O., Pincet, F., Perez, E., Thiery, J.P., and Dufour, S. (2004). Force measurements in E-cadherin-mediated cell doublets reveal rapid adhesion strengthened by actin cytoskeleton remodeling through Rac and Cdc42. J. Cell Biol. 167, 1183–1194.

17. Daniel, E., Daudé, M., Kolotuev, I., Charish, K., Auld, V., and Le Borgne, R. (2018). Coordination of Septate Junctions Assembly and Completion of Cytokinesis in Proliferative Epithelial Tissues. Curr. Biol. 28, 1380–1391.e4.

18. Davis, J.R., Ainslie, A.P., Williamson, J.J., Ferreira, A., Torres-Sánchez, A., Hoppe, A., Mangione, F., Smith, M.B., Martin-Blanco, E., Salbreux, G., et al. (2022). ECM degradation in the Drosophila abdominal epidermis initiates tissue growth that ceases with rapid cell-cycle exit. Curr. Biol. 32, 1285–1300.e4.

19. Delague, V., Jacquier, A., Hamadouche, T., Poitelon, Y., Baudot, C., Boccaccio, I., Chouery, E., Chaouch, M., Kassouri, N., Jabbour, R., et al. (2007). Mutations in FGD4 encoding the Rho GDP/GTP exchange factor FRABIN cause autosomal recessive Charcot-Marie-Tooth type 4H. Am. J. Hum. Genet. 81, 1–16.

20. Denk-Lobnig, M., and Martin, A.C. (2019). Modular regulation of Rho family GTPases in development. Small GTPases.

21. Dent, L.G., Manning, S.A., Kroeger, B., Williams, A.M., Hilmi, A.J.S., Crea, L., Kondo, S., Badovinac, S.H., and Harvey, K.F. (2019). The dPix-Git complex is essential to coordinate epithelial morphogenesis and regulate myosin during Drosophila egg chamber development. PLoS Genet. 15, e1008083.

22. Duhart, J.C., Parsons, T.T., and Raftery, L.A. (2017). The repertoire of epithelial morphogenesis on display: Progressive elaboration of Drosophila egg structure. Mech. Dev. 148, 18–39.

23. Duszyc, K., Gomez, G.A., Lagendijk, A.K., Yau, M.K., Nanavati, B.N., Gliddon, B.L., Hall, T.E., Verma, S., Hogan, B.M., Pitson, S.M., et al. (2021). Mechanotransduction activates RhoA in the neighbors of apoptotic epithelial cells to engage apical extrusion. Curr. Biol. 31, 1326–1336.e5.

24. Fernandez-Gonzalez, R., and Peifer, M. (2022). Powering morphogenesis: multiscale challenges at the interface of cell adhesion and the cytoskeleton. Mol. Biol. Cell 33.

25. Fic, W., Bastock, R., Raimondi, F., Los, E., Inoue, Y., Gallop, J.L., Russell, R.B., and Johnston, D.S. (2021). RhoGAP19D inhibits Cdc42 laterally to control epithelial cell shape and prevent invasion. J. Cell Biol. 220, e202009116.

26. Firmino, J., Rocancourt, D., Saadaoui, M., Moreau, C., and Gros, J. (2016). Cell Division Drives Epithelial Cell Rearrangements during Gastrulation in Chick. Dev. Cell 36, 249–261.

27. Founounou, N., Loyer, N., and Le Borgne, R. (2013). Septins Regulate the Contractility of the Actomyosin Ring to Enable Adherens Junction Remodeling during Cytokinesis of Epithelial Cells. Dev. Cell 24, 242–255.

28. Garcia De Las Bayonas, A., Philippe, J.M., Lellouch, A.C., and Lecuit, T. (2019). Distinct RhoGEFs Activate Apical and Junctional Contractility under Control of G Proteins during Epithelial Morphogenesis. Curr. Biol. 29, 3370–3385.e7.

29. Georgiou, M., and Baum, B. (2010). Polarity proteins and Rho GTPases cooperate to spatially organise epithelial actin-based protrusions. J. Cell Sci. 123, 1089–1098.

30. Gibson, M.C., Patel, A.B., Nagpal, R., and Perrimon, N. (2006). The emergence of geometric order in proliferating metazoan epithelia. Nature 442, 1038–1041.

31. Glotzer, M. (2017). Cytokinesis in metazoa and fungi. Cold Spring Harb. Perspect. Biol. 9, a022343.

32. Godard, B.G., and Heisenberg, C.P. (2019). Cell division and tissue mechanics. Curr. Opin. Cell Biol. 60, 114–120.

33. Golic, K.G., and Lindquist, S. (1989). The FLP recombinase of yeast catalyzes site-specific recombination in the drosophila genome. Cell 59, 499–509.

34. Gratz, S.J., Ukken, F.P., Rubinstein, C.D., Thiede, G., Donohue, L.K., Cummings, A.M., and Oconnor-Giles, K.M. (2014). Highly specific and efficient CRISPR/Cas9-catalyzed homology- directed repair in Drosophila. Genetics 196, 961–971.

35. Guillot, C., and Lecuit, T. (2013). Adhesion Disengagement Uncouples Intrinsic and Extrinsic Forces to Drive Cytokinesis in Epithelial Tissues. Dev. Cell 24, 227–241.

36. Guirao, B., Rigaud, S.U., Bosveld, F., Bailles, A., López-Gay, J., Ishihara, S., Sugimura, K., Graner, F., and Bellaïche, Y. (2015). Unified quantitative characterization of epithelial tissue development. Elife 4, e08519.

37. Heasman, S.J., and Ridley, A.J. (2008). Mammalian Rho GTPases: new insights into their functions from in vivo studies. Nat. Rev. Mol. Cell Biol. 9, 690–701.

38. Heller, D., Hoppe, A., Restrepo, S., Gatti, L., Tournier, A.L., Tapon, N., Basler, K., and Mao, Y. (2016). EpiTools: An Open-Source Image Analysis Toolkit for Quantifying Epithelial Growth Dynamics. Dev. Cell 36, 103–116.

39. Herszterg, S., Leibfried, A., Bosveld, F., Martin, C., and Bellaiche, Y. (2013). Interplay between the Dividing Cell and Its Neighbors Regulates Adherens Junction Formation during Cytokinesis in Epithelial Tissue. Dev. Cell 24, 256–270.

40. Herszterg, S., Pinheiro, D., and Bellaïche, Y. (2014). A multicellular view of cytokinesis in epithelial tissue. Trends Cell Biol. 24, 285–293.

41. Higashi, T., Arnold, T.R., Stephenson, R.E., Dinshaw, K.M., and Miller, A.L. (2016). Maintenance of the Epithelial Barrier and Remodeling of Cell-Cell Junctions during Cytokinesis. Curr. Biol. 26, 1829–1842.

42. Hodge, R.G., and Ridley, A.J. (2016). Regulating Rho GTPases and their regulators. Nat. Rev. Mol. Cell Biol. 17, 496–510.

43. Huang, J., Zhou, W., Watson, A.M., Jan, Y.N., and Hong, Y. (2008). Efficient ends-out gene targeting in drosophila. Genetics 180, 703–707.

44. Iden, S., and Collard, J.G. (2008). Crosstalk between small GTPases and polarity proteins in cell polarization. Nat. Rev. Mol. Cell Biol. 9, 846–859.

45. Jaffe, A.B., and Hall, A. (2005). Rho GTPases: biochemistry and biology. Annu. Rev. Cell Dev. Biol. 21, 247–269.

46. Jones, W.M., Chao, A.T., Zavortink, M., Saint, R., and Bejsovec, A. (2010). Cytokinesis proteins Tum and Pav have a nuclear role in Wnt regulation. J. Cell Sci. 123, 2179–2189.

47. Jordan, S.N., and Canman, J.C. (2012). Rho GTPases in animal cell cytokinesis: An occupation by the one percent. Cytoskeleton 69, 919–930.

48. Jülicher, F., and Eaton, S. (2017). Emergence of tissue shape changes from collective cell behaviours. Semin. Cell Dev. Biol. 67, 103–112.

49. Kanca, O., Zirin, J., Garcia-Marques, J., Knight, S.M., Yang-Zhou, D., Amador, G., Chung, H., Zuo, Z., Ma, L., He, Y., et al. (2019). An efficient CRISPR-based strategy to insert small and large fragments of DNA using short homology arms. Elife 8, e51539.

50. Lau, K., Tao, H., Liu, H., Wen, J., Sturgeon, K., Sorfazlian, N., Lazic, S., Burrows, J.T.A.A., Wong, M.D., Li, D., et al. (2015). Anisotropic stress orients remodelling of mammalian limb bud ectoderm. Nat. Cell Biol. 17, 569–579.

51. Laurin, M., Gomez, N.C., Levorse, J., Sendoel, A., Sribour, M., and Fuchs, E. (2019). An RNAi screen unravels the complexities of rho GTPase networks in skin morphogenesis. Elife 8, 1–34.

52. Lawson, C.D., and Ridley, A.J. (2018). Rho GTPase signaling complexes in cell migration and invasion. J. Cell Biol. 217, 447–457.

53. Lechler, T., and Mapelli, M. (2021). Spindle positioning and its impact on vertebrate tissue architecture and cell fate. Nat. Rev. Mol. Cell Biol. 22, 691–708.

54. Lecuit, T., and Yap, A.S. (2015). E-cadherin junctions as active mechanical integrators in tissue dynamics. Nat. Cell Biol. 17, 533–539.

55. Lee, S., Park, H., Kyung, T., Kim, N.Y., Kim, S., Kim, J., and Heo, W. Do (2014). Reversible protein inactivation by optogenetic trapping in cells. Nat. Methods 11, 633–636.

56. Li, M.Z., and Elledge, S.J. (2012). SLIC: A method for sequence- and ligation-independent cloning. Methods Mol. Biol. 852, 51–59.

57. Loria, A., Longhini, K.M., and Glotzer, M. (2012). The RhoGAP domain of CYK-4 Has an essential role in RhoA activation. Curr. Biol. 22, 213–219.

58. Mason, F.M., Xie, S., Vasquez, C.G., Tworoger, M., and Martin, A.C. (2016). RhoA GTPase inhibition organizes contraction during epithelial morphogenesis. J. Cell Biol. 214, 603–617.

59. McGuire, S.E., Le, P.T., Osborn, A.J., Matsumoto, K., and Davis, R.L. (2003). Spatiotemporal Rescue of Memory Dysfunction in Drosophila. Science (80-.). 302, 1765–1768.

60. McKinley, K.L., Stuurman, N., Royer, L.A., Schartner, C., Castillo-Azofeifa, D., Delling, M., Klein, O.D., and Vale, R.D. (2018). Cellular aspect ratio and cell division mechanics underlie the patterning of cell progeny in diverse mammalian epithelia. Elife 7, e36739.

61. Mishima, M., Kaitna, S., and Glotzer, M. (2002). Central spindle assembly and cytokinesis require a kinesin-like protein/RhoGAP complex with microtubule bundling activity. Dev. Cell 2, 41–54.

62. Monster, J.L., Donker, L., Vliem, M.J., Win, Z., Matthews, H.K., Cheah, J.S., Yamada, S., de Rooij, J., Baum, B., and Gloerich, M. (2021). An asymmetric junctional mechanoresponse coordinates mitotic rounding with epithelial integrity. J. Cell Biol. 220, e202001042.

63. Morais-De-Sá, E., and Sunkel, C. (2013). Adherens junctions determine the apical position of the midbody during follicular epithelial cell division. EMBO Rep. 14, 696–703.

64. Müller, P.M., Rademacher, J., Bagshaw, R.D., Wortmann, C., Barth, C., van Unen, J., Alp, K.M., Giudice, G., Eccles, R.L., Heinrich, L.E., et al. (2020). Systems analysis of RhoGEF and RhoGAP regulatory proteins reveals spatially organized RAC1 signalling from integrin adhesions. Nat. Cell Biol. 22, 498–511.

65. Munjal, A., Philippe, J.M., Munro, E., and Lecuit, T. (2015). A self-organized biomechanical network drives shape changes during tissue morphogenesis. Nature 524, 351–355.

66. Nahm, M., Lee, M., Baek, S.H., Yoon, J.H., Kim, H.H., Lee, Z.H., and Lee, S. (2006). Drosophila RhoGEF4 encodes a novel RhoA-specific guanine exchange factor that is highly expressed in the embryonic central nervous system. Gene 384, 139–144.

67. Nakajima, H., and Tanoue, T. (2011). Lulu2 regulates the circumferential actomyosin tensile system in epithelial cells through p114RhoGEF. J. Cell Biol. 195, 245–261.

68. Neisch, A.L., Formstecher, E., and Fehon, R.G. (2013). Conundrum, an ARHGAP18 orthologue, regulates RhoA and proliferation through interactions with Moesin. Mol. Biol. Cell 24, 1420–1433.

69. Ninov, N., Manjón, C., and Martín-Blanco, E. (2009). Dynamic control of cell cycle and growth coupling by ecdysone, egfr, and PI3K signaling in Drosophila histoblasts. PLoS Biol. 7, e1000079.

70. Olmos, Y., Perdrix-Rosell, A., and Carlton, J.G. (2016). Membrane Binding by CHMP7 Coordinates ESCRT-III-Dependent Nuclear Envelope Reformation. Curr. Biol. 26, 2635–2641.

71. Osswald, M., Barros-Carvalho, A., Carmo, A.M., Loyer, N., Gracio, P.C., Sunkel, C.E., Homem, C.C.F., Januschke, J., and Morais-de-Sá, E. (2022). aPKC regulates apical constriction to prevent tissue rupture in the Drosophila follicular epithelium. Curr. Biol. S0960- 9822(22)01385-9.

72. Pinheiro, D., and Bellaïche, Y. (2018). Mechanical Force-Driven Adherens Junction Remodeling and Epithelial Dynamics. Dev. Cell 47, 3–19.

73. Pinheiro, D., Hannezo, E., Herszterg, S., Bosveld, F., Gaugue, I., Balakireva, M., Wang, Z., Cristo, I., Rigaud, S.U., Markova, O., et al. (2017). Transmission of cytokinesis forces via E- cadherin dilution and actomyosin flows. Nature 545, 103–107.

74. Port, F., Chen, H.-M., Lee, T., and Bullock, S.L. (2014). Optimized CRISPR/Cas tools for efficient germline and somatic genome engineering in Drosophila. Proc. Natl. Acad. Sci. U. S. A. 111, E2967–76.

75. Prokopenko, S.N., Saint, R., and Bellen, H.J. (2000). Tissue distribution of PEBBLE RNA and pebble protein during Drosophila embryonic development. Mech. Dev. 90, 269–273.

76. Qin, X., Park, B.O., Liu, J., Chen, B., Choesmel-Cadamuro, V., Belguise, K., Heo, W. Do, and Wang, X. (2017). Cell-matrix adhesion and cell-cell adhesion differentially control basal myosin oscillation and Drosophila egg chamber elongation. Nat. Commun. 8, 14708.

77. Ragkousi, K., and Gibson, M.C. (2014). Cell division and the maintenance of epithelial order. J. Cell Biol. 207, 181–188.

78. Ramkumar, N., and Baum, B. (2016). Coupling changes in cell shape to chromosome segregation. Nat. Rev. Mol. Cell Biol. 17, 511–521.

79. Renda, I., Bianchi, S., Vezzosi, V., Nori, J., Vanzi, E., Tavella, K., and Susini, T. (2019). Expression of FGD3 gene as prognostic factor in young breast cancer patients. Sci. Rep. 9, 15204.

80. Rosa, A., Vlassaks, E., Pichaud, F., and Baum, B. (2015). Ect2/Pbl acts via Rho and polarity proteins to direct the assembly of an isotropic actomyosin cortex upon mitotic entry. Dev. Cell 32, 604–616.

81. Schmidt, A., Lv, Z., and Großhans, J. (2018). Elmo and sponge specify subapical restriction of canoe and formation of the subapical domain in early drosophila embryos. Dev. 145, dev157909.

82. Silver, J.T., Wirtz-Peitz, F., Simões, S., Pellikka, M., Yan, D., Binari, R., Nishimura, T., Li, Y., Harris, T.J.C., Perrimon, N., et al. (2019). Apical polarity proteins recruit the RhoGEF Cysts to promote junctional myosin assembly. J. Cell Biol. 218, 3397–3414.

83. Simões, S., Lerchbaumer, G., Pellikka, M., Giannatou, P., Lam, T., Kim, D., Yu, J., Stal, D. Ter, Al Kakouni, K., Fernandez-Gonzalez, R., et al. (2022). Crumbs complex–directed apical membrane dynamics in epithelial cell ingression. J. Cell Biol. 221, e202108076.

84. Su, K.C., Takaki, T., and Petronczki, M. (2011). Targeting of the RhoGEF Ect2 to the Equatorial Membrane Controls Cleavage Furrow Formation during Cytokinesis. Dev. Cell 21, 1104–1115.

85. Sugimura, K., and Ishihara, S. (2013). The mechanical anisotropy in a tissue promotes ordering in hexagonal cell packing. Development 140, 4091–4101.

86. Taubenberger, A. V., Baum, B., and Matthews, H.K. (2020). The Mechanics of Mitotic Cell Rounding. Front. Cell Dev. Biol. 8, 687.

87. Toret, C.P., Collins, C., and Nelson, W.J. (2014). An Elmo-Dock complex locally controls Rho GTPases and actin remodeling during cadherin-mediated adhesion. J. Cell Biol. 207, 577–587.

88. Toret, C.P., Shivakumar, P.C., Lenne, P.F., and Le Bivic, A. (2018). The ELMO-MBC complex and RhoGAP19D couple rho family GTPases during mesenchymal-to-epithelial-like transitions. Dev. 145, dev.157495.

89. Uroz, M., Wistorf, S., Serra-Picamal, X., Conte, V., Sales-Pardo, M., Roca-Cusachs, P., Guimerà, R., and Trepat, X. (2018). Regulation of cell cycle progression by cell–cell and cell– matrix forces. Nat. Cell Biol. 20, 646–654.

90. Venken, K.J.T., Kasprowicz, J., Kuenen, S., Yan, J., Hassan, B.A., and Verstreken, P. (2008). Recombineering-mediated tagging of Drosophila genomic constructs for in vivo localization and acute protein inactivation. 36, 1–9.

91. Venken, K.J.T., Carlson, J.W., Schulze, K.L., Pan, H., He, Y., Spokony, R., Wan, K.H., Koriabine, M., de Jong, P.J., White, K.P., et al. (2009). Versatile P[acman] BAC libraries for transgenesis studies in Drosophila melanogaster. Nat. Methods 6, 431–434.

92. Verma, S., Han, S.P., Michael, M., Gomez, G.A., Yang, Z., Teasdale, R.D., Ratheesh, A., Kovacs, E.M., Ali, R.G., and Yap, A.S. (2012). A WAVE2-Arp2/3 actin nucleator apparatus supports junctional tension at the epithelial zonula adherens. Mol. Biol. Cell 23, 4601–4610.

93. Wang, Z., Bosveld, F., and Bellaïche, Y. (2018). Tricellular junction proteins promote disentanglement of daughter and neighbour cells during epithelial cytokinesis. J. Cell Sci. 131, jcs215764.

94. Yamada, S., and Nelson, W.J. (2007). Localized zones of Rho and Rac activities drive initiation and expansion of epithelial cell-cell adhesion. J. Cell Biol. 178, 517–527.

95. Yüce, Ö., Piekny, A., and Glotzer, M. (2005). An ECT2-centralspindlin complex regulates the localization and function of RhoA. J. Cell Biol. 170, 571–582.

96. Zhang, X., Koolhaas, W.H., and Schnorrer, F. (2014). A versatile two-step CRISPR- and RMCE-based strategy for efficient genome engineering in Drosophila. G3 Genes, Genomes, Genet. 4, 2409–2418.

97. Zhao, W.M., and Fang, G. (2005). MgcRacGAP controls the assembly of the contractile ring and the initiation of cytokinesis. Proc. Natl. Acad. Sci. U. S. A. 102, 13158–13163.

98. Zihni, C., Munro, P.M.G., Elbediwy, A., Keep, N.H., Terry, S.J., Harris, J., Balda, M.S., and Matter, K. (2014). Dbl3 drives Cdc42 signaling at the apical margin to regulate junction position and apical differentiation. J. Cell Biol. 204, 111–127.

