## Supplementary material for "Systematic characterization of *Drosophila* RhoGEF/GAP localizations uncovers regulators of mechanosensing and junction formation during epithelial cell division": Key Ressource Table

**Key resources table**

| REAGENT or RESOURCE | SOURCE | IDENTIFIER |
|---|---|---|
| Experimental models: Organisms/strains | | |
| Drosophila: Act5C-Gal4 | Bloomington Drosophila Stock Center. | BDSC#3954 |
| Drosophila: tub-Gal80$^{ts}$ | (McGuire et al., 2001) ; Bloomington Drosophila Stock Center | BDSC#7017 |
| Drosophila : Act5c>cd2>GAL4 | Bloomington Drosophila Stock Center | BDSC#4780 |
| Drosophila: hs-flp | Bloomington Drosophila Stock Center | |
| Drosophila: FRT80B, nls:GFP | Bloomington Drosophila Stock Center | BDSC #5630 |
| Drosophila: FRT80B, H2A:RFP | Bloomington Drosophila Stock Center | BDSC#34499 |
| Drosophila: FRT40A, nls:GFP | Bloomington Drosophila Stock Center | BDSC#5629 |
| Drosophila: FRT40A, ubi-H2B:RFP | (Bosveld et al., 2017) | NA |
| Drosophila:Cyo,hs-Cre | Bloomington Drosophila Stock Center | BDSC#1092 |
| Drosophila:TM6,hs-Cre | Bloomington Drosophila Stock Center | BDSC#1501 |
| Drosophila: Ecad:GFP$^{ki}$ | (Huang et al., 2009) ; gift of Yang Hong | NA |
| Drosophila: Ecad:3xmKate2 | (Pinheiro et al., 2017) | NA |
| Drosophila: sqh:3xmKate2 | (Pinheiro et al., 2017) | NA |
| Drosophila: sqh:3xGFP | (Pinheiro et al., 2017) | NA |
| Drosophila: UAS-CAAX:tBFP (II) | (López-Gay et al., 2020) | NA |
| Drosophila: UAS-CAAX:tBFP (III) | (López-Gay et al., 2020) | NA |
| Drosophila: UAS-LARIAT | (Qin et al., 2017) | NA |
| Drosophila: sqh-Utr:ABD:GFP | (Rauzi et al., 2010) | NA |
| Drosophila: ubi-PLCgPH:GFP | Gift from F. Pichaud | NA |
| Drosophila: ubi-PLCgPH:chFP | (Herszterg et al., 2013) | NA |
| Drosophila: ubi-Anillin-Rho binding Domain:GFP | (Munjal et al., 2015), gift from T. Lecuit | NA |
| Drosophila: Rac1:GFP | Bloomington Drosophila Stock Center | BDSC#52284 |
| Drosophila: sqh-Pak3-RhoGTPases Binding Domain: GFP | Bloomington Drosophila Stock Center | BDSC#52303 |
| Drosophila: Scar:GFP | This study | NA |
| Drosophila: Lamin:TagRFP$^{ki}$ | (Ambrosini et al., 2019), gift from M.Suzanne | NA |
| Drosophila: Nup107:RFP | Bloomington Drosophila Stock Center | BDSC#35517 |
| Drosophila: ubi-Spd2:RFP | (Conduit et al., 2014) | NA |
| Drosophila: *rhogef4*$^{\square RMCE}$ | This study | NA |
| Drosophila: *cysts*$^{\square RMCE}$ | This study | NA |
| Drosophila: FRT *Rho1*$^{72M1}$ | (Strutt et al., 1997) | NA |
| Drosophila: Df (3L) BSC117 | Bloomington Drosophila Stock Center | BDSC#8974 |
| Drosophila: UAS-pnut$^{dsRNA}$ | VDRC Stock Center, (Dietzl et al., 2007) | VDRC#11791 |
| Drosophila: UAS-shg$^{dsRNA}$ | VDRC Stock Center, (Dietzl et al., 2007) | VDRC#27082 |
| Drosophila : UAS-Rok$^{dsRNA}$ | Bloomington Drosophila Stock Center | BDSC28797 |
| Drosophila: vas-cas9 | Bloomington Drosophila Stock Center. | BDSC#55821 |
| Drosophila: vas-cas9 | Bloomington Drosophila Stock Center. | BDSC#51324 |

| | | |
|---|---|---|
| Drosophila: fluorescently tagged RhoGEF/ GAP | This study, see Supplementary Table S1. | |
| Oligonucleotides | | |
| See Table S1 | | |
| Recombinant DNA | | |
| See Table S1 | | |
| Chemicals | | |
| Insulin solution from bovine pancreas | Sigma-Aldrich | Cat# I0516 |
| Schneider's Insect Medium | Sigma-Aldrich | Cat# S0146 |
| Cell Mask Orange Plasma membrane Stain | ThermoFisher | Cat# C10045 |
| Software and algorithms | | |
| Fiji | http://fiji.sc | SCR_002285 |
| MetaMorph Microscopy Automation and Image Analysis Software | Molecular devices http://www.moleculardevices.com/Products/Software/Meta-Imaging-Series/MetaMorph.html | SCR_002368 |
| ZEN Digital Imaging for Light Microscopy | Zeiss http://www.zeiss.com/microscopy/en_us/products/microscope-software/zen.html#introduction | SCR_013672 |
| Graphpad Prism | Graphpad software http://www.graphpad.com/ | SCR_002798 |
| Matlab | Mathworks http://www.mathworks.com/products/matlab/ | SCR_001622 |

**References:**


Ambrosini, A., Rayer, M., Monier, B., and Suzanne, M. (2019). Mechanical Function of the Nucleus in Force Generation during Epithelial Morphogenesis. Dev. Cell *50*, 197-211.e5.

Bosveld, F., Ainslie, A., and Bellaïche, Y. (2017). Sequential activities of Dynein, Mud and Asp in centrosome- spindle coupling maintain centrosome number upon mitosis. J. Cell Sci. *130*, 3557–3567.

Conduit, P.T., Feng, Z., Richens, J.H., Baumbach, J., Wainman, A., Bakshi, S.D., Dobbelaere, J., Johnson, S., Lea, S.M., and Raff, J.W. (2014). The centrosome-specific phosphorylation of Cnn by Polo/Plk1 drives Cnn scaffold assembly and centrosome maturation. Dev. Cell *28*, 659–669.

Dietzl, G., Chen, D., Schnorrer, F., Su, K.C., Barinova, Y., Fellner, M., Gasser, B., Kinsey, K., Oppel, S., Scheiblauer, S., et al. (2007). A genome-wide transgenic RNAi library for conditional gene inactivation in Drosophila. Nature *448*, 151–156.

Herszterg, S., Leibfried, A., Bosveld, F., Martin, C., and Bellaiche, Y. (2013). Interplay between the Dividing Cell and Its Neighbors Regulates Adherens Junction Formation during Cytokinesis in Epithelial Tissue. Dev. Cell *24*, 256–270.

Huang, J., Zhou, W., Dong, W., Watson, A.M., and Hong, Y. (2009). From the Cover: Directed, efficient, and versatile modifications of the Drosophila genome by genomic engineering. Proc. Natl. Acad. Sci. U. S. A. *106*, 8284–8289.

López-Gay, J.M., Nunley, H., Spencer, M., di Pietro, F., Guirao, B., Bosveld, F., Markova, O., Gaugue, I., Pelletier, S., Lubensky, D.K., et al. (2020). Apical stress fibers enable a scaling between cell mechanical response and area in epithelial tissue. Science (80-. ). *370*.

McGuire, S.E., Le, P.T., and Davis, R.L. (2001). The role of Drosophila mushroom body signaling in olfactory memory. Science (80-. ). *293*, 1330–1333.

Munjal, A., Philippe, J.M., Munro, E., and Lecuit, T. (2015). A self-organized biomechanical network drives shape changes during tissue morphogenesis. Nature *524*, 351–355.

Pinheiro, D., Hannezo, E., Herszterg, S., Bosveld, F., Gaugue, I., Balakireva, M., Wang, Z., Cristo, I., Rigaud, S.U., Markova, O., et al. (2017). Transmission of cytokinesis forces via E-cadherin dilution and actomyosin flows. Nature *545*, 103–107.

Qin, X., Park, B.O., Liu, J., Chen, B., Choesmel-Cadamuro, V., Belguise, K., Heo, W. Do, and Wang, X. (2017). Cell-matrix adhesion and cell-cell adhesion differentially control basal myosin oscillation and Drosophila egg chamber elongation. Nat. Commun. *8*, 14708.

Rauzi, M., Lenne, P., and Lecuit, T. (2010). Planar polarized actomyosin contractile flows control epithelial junction remodelling. Nature *468*, 1110–1114.

Strutt, D.I., Weber, U., and Mlodzik, M. (1997). The role of RhoA in tissue polarity and Frizzled signalling. Nature *387*, 292–295.
